# Tumor cell-based liquid biopsy using high-throughput microfluidic enrichment of entire leukapheresis product

**DOI:** 10.1101/2024.03.13.583573

**Authors:** Avanish Mishra, Shih-Bo Huang, Taronish Dubash, Risa Burr, Jon F. Edd, Ben S. Wittner, Quinn E. Cunneely, Victor R. Putaturo, Akansha Deshpande, Ezgi Antmen, Kaustav A. Gopinathan, Keisuke Otani, Yoshiyuki Miyazawa, Ji Eun Kwak, Sara Y. Guay, Justin Kelly, John Walsh, Linda Nieman, Isabella Galler, PuiYee Chan, Michael S. Lawrence, Ryan J. Sullivan, Aditya Bardia, Douglas S. Micalizzi, Lecia V. Sequist, Richard J. Lee, Joseph W. Franses, David T. Ting, Patricia A. R. Brunker, Shyamala Maheswaran, David T. Miyamoto, Daniel A. Haber, Mehmet Toner

## Abstract

Circulating Tumor Cells (CTCs), interrogated by sampling blood from patients with cancer, contain multiple analytes, including intact RNA, high molecular weight DNA, proteins, and metabolic markers. However, the clinical utility of tumor cell-based liquid biopsy has been limited since CTCs are very rare, and current technologies cannot process the blood volumes required to isolate a sufficient number of tumor cells for in-depth assays. We previously described a high-throughput microfluidic prototype utilizing high-flow channels and amplification of cell sorting forces through magnetic lenses. Here, we apply this technology to analyze patient-derived leukapheresis products, interrogating a mean blood volume of 5.83 liters from patients with metastatic cancer, with a median of 2,799 CTCs purified per patient. Isolation of many CTCs from individual patients enables characterization of their morphological and molecular heterogeneity, including cell and nuclear size and RNA expression. It also allows robust detection of gene copy number variation, a definitive cancer marker with potential diagnostic applications. High-volume microfluidic enrichment of CTCs constitutes a new dimension in liquid biopsies.

## Main

Liquid biopsies provide increasingly important non-invasive strategies for longitudinal monitoring of cancer, guiding personalized therapies as tumor cells evolve under selective pressures^1^. DNA sequencing-based analyses of tumor-derived circulating DNA fragments (ctDNA) enable the detection of drug-sensitizing and drug-resistant genomic mutations^2^. ctDNA-based approaches are now being tested to detect minimal residual disease following surgery^3^ and early recurrence based on mutational or DNA methylation abnormalities^4^. Intact circulating tumor cells (CTCs) are also shed into the bloodstream, and while they provide the full complement of analytes, including high molecular weight DNA, intact RNA, cytoplasmic and cell surface proteins, and metabolic markers, the rarity of CTCs in the blood has limited their clinical utility^5,6^.

A standard 10 mL blood tube drawn from the peripheral circulation may yield 0 to 10 CTCs, depending on tumor histology and stage, as well as the type of assay applied to enrich CTCs^6,7^. Most studies report the absence of any detectable CTCs in 20-50% of patients^8,9^. Even when CTCs are readily detected within 10 mL of blood, their numbers may be too low to allow for statistically robust analytics, given the heterogeneity in the expression of cancer-relevant markers and the analytical sensitivity limits posed by most assays. Nonetheless, the emergence of highly effective antibody-based therapies directed against tumor epitopes highlights the critical unmet need for reliable cell-based liquid biopsies with quantitation of specific protein expression^10,11^. Needle biopsies of accessible individual metastatic lesions during the course of treatment have proven to be an important strategy to tailor therapeutic choices, but these are not readily repeated serially due to the invasive nature of this procedure^12^. Moreover, they sample a single site of disease, which does not capture the heterogeneity of metastatic cancer^13^. Given the rapid turnover of CTCs and the potential representation of all invasive tumor deposits within a single specimen, a blood-based technology that can generate sufficient numbers of intact cancer cells for reliable analysis would be highly impactful in guiding drug development and clinical treatment choices.

Leukapheresis is an established method for isolating rare cell populations within the blood, including hematopoietic stem cells for bone marrow transplantation or large numbers of T cells for CAR-T cell engineering^14^. In this standard clinical procedure, typically performed at a blood bank or apheresis center, blood is drawn from the antecubital vein in one arm and centrifuged through a continuous process that separates mononuclear cells from red blood cells (RBCs), plasma, and platelets, which are then returned to the patient through the contralateral antecubital vein^15–18^. The sedimentation of CTCs overlaps with that of mononuclear leukocytes, making it possible to enrich these cells using standard leukapheresis parameters^17^. A typical leukapheresis procedure interrogates 3L of blood volume per hour^19^. Due to their rapid turnover, removed white blood cells (WBCs) and platelets are rapidly regenerated by the patient, and leukapheresis is generally associated with less than 20 mL of net RBC loss. The primary clinical criteria for tolerating leukapheresis include adequate venous access and cardiovascular stability, given the need for anticoagulation and the possibility of transient blood pressure and intravascular volume shifts^18,19^.

To realize the promise of using leukapheresis to isolate sufficient numbers of CTCs in patients with cancer, the complexities of rare cell enrichment technologies need to be addressed. The leukapheresis product (leukopak) contains an extremely high total number of WBCs and platelets (50 to 100-fold higher than whole blood), within a large sample volume of approximately 100 mL.

The FDA-approved CellSearch technology, which uses antibody-bound magnetic ferrofluids to enrich for CTCs expressing the epithelial cell surface marker EpCAM, can only process 5% of a leukopak^15,17^. Initial applications of this technology to leukopak samples have confirmed the expected increase in CTC capture, with an 11.5-fold increase in CTCs obtained from 5% leukopak (approximately 2% total blood volume), compared with the standard CellSearch CTC yield from a 7.5mL blood tube (approximately 0.15% total blood volume)^15^. We previously reported a microfluidic CTC enrichment technology that does not require cell fixation and is sufficiently efficient to allow 10^4^-fold depletion of WBCs using magnetized antibodies, thereby enriching CTCs independently of individual tumor epitopes^7,20^. This CTC-iChip “negative depletion” technology can process 20mL of whole blood within one hour, but like other microfluidic technologies, it ultimately suffers from clogging and reduced throughput as larger and more concentrated fluids are processed^20,21^. To address these challenges and enable the sorting of the substantial numbers of blood cells present in a leukopak, we recently created an ultra-high-throughput microfluidic platform (^LP^CTC-iChip)^22^. In this platform technology, initial microfluidic debulking of RBCs and platelets is applied, followed by flowing nucleated blood cells through “magnetic lenses” made of magnetized microchannels positioned within 60 µm of the blood channels, thereby enhancing magnetic forces by 35-fold and enhancing sorting throughput by 30-fold. The engineering optimization required for the ^LP^CTC-iChip and its efficacy in capturing 86.1±0.6% of individual cancer cells spiked into control healthy donor leukopaks were previously described^22^. Here, we describe the first application of the ^LP^CTC-iChip to processing entire leukopaks (1-5L of blood volume equivalent) in patients with metastatic cancer and the initial cellular and molecular characterization of the unprecedentedly large numbers of CTCs from individual patients.

## Results

### Clinical cases

Six patients with metastatic cancer undergoing treatment at the Mass General Cancer Center consented to leukapheresis for research purposes (MGB IRB 2020P000251). Leukapheresis was performed at the Mass General Blood Transfusion Service department. All patients tolerated the procedure without any reported adverse events. Leukopaks were maintained at room temperature and processed within 6 hours of collection. Table I presents a brief clinical history of each patient.

**Table 1:** Clinical characteristics of patients.

| Patient | Diagnosis | Age | Gender | Years Since Primary Diagnosis | Sites of Disease | Disease Status at Time of Leukapheresis | Treatment History |
| --- | --- | --- | --- | --- | --- | --- | --- |
| **GU-1** | Metastatic Prostate Cancer | 69 | M | 19 | Bones, lymph nodes, adrenal glands, and peritoneum | Progressive disease | <u>Localized</u>: RT and ADT<br><u>Metastatic</u>: ADT, darolutamide, abiraterone, cabozantinib/atezolizumab, docetaxel, cabazitaxel, lutetium-177 vipivotide tetraxetan |
| **TNBC-1** | Metastatic Triple Negative Breast Cancer | 35 | F | 5 | Lungs, lymph nodes, bone, and liver | Stable disease | <u>Localized</u>: Neoadjuvant paclitaxel/fluorouracil/doxorubicin/cyclophosphamide, surgery, and RT<br><u>Metastatic</u>: paclitaxel/carboplatin/atezolizumab, capecitabine/bevacizumab, sacituzumab govitecan, and Talazoparib |
| **HCC-1** | Metastatic Hepatocellular carcinoma | 60 | M | 4 | Liver, retrocaval lymph nodes, lungs | Stable disease | <u>Localized</u>: gelfoam embolization<br><u>Metastatic</u>: liver embolization/RT, atezolizumab/bevacizumab |
| **HCC-2** | Metastatic Hepatocellular carcinoma | 70 | M | 3 | Lung, bone | Stable disease | <u>Localized</u>: RT<br><u>Metastatic</u>: atezolizumab/bevacizumab, regorafenib/pembrolizumab |
| **UM-1** | Metastatic Uveal Melanoma | 75 | M | 4 | Liver, bone, lung, soft tissue | Stable disease | <u>Localized</u>: RT<br><u>Metastatic</u>: pembrolizumab, yttrium-90 irradiation of liver, tebentafusp, |
| **UM-2** | Metastatic Uveal Melanoma | 55 | M | 2 | Liver, bone, lung, soft tissue | Progressive disease | <u>Localized</u>: RT<br><u>Metastatic</u>: tebentafusp, nivolumab/ipilimumab |

Abbreviations: ADT, androgen deprivation therapy and RT, radiation therapy

### Parameters for leukapheresis and leukopak characterization

The strategy for isolating CTCs from leukapheresis samples collected from patients with cancer is schematically illustrated in **Fig. 1**. Leukopaks were generated from six patients with metastatic cancer, interrogating approximately one full blood volume (5.83 ± 1.03 L) over a 2-hour period in continuous mononuclear cell collection mode (Spectra Optia) (see Methods for leukapheresis settings). By virtue of their comparable sedimentation rate, CTCs are collected in the same layers as mononuclear leukocytes^14,15^. Leukapheresis was performed at a flow rate of 53.6 ± 7.4 mL/min, with the mean volume of collected leukopaks 107.7±4.7 mL (**Fig. 2A-B).** The mean WBC concentration of leukopaks was 49.4 ± 23.7×10^6^ cells/mL (range 21.1×10^6^ cells/mL to 95.8×10^6^ cells/mL), which corresponds to an approximately 8-fold higher concentration than that of normal whole blood. The inter-sample variability in WBC concentration was as high as 4-fold, which is considerably higher than the variability observed in whole blood samples. The mean platelet concentration of leukopaks was 1.4 ± 0.4×10^9^ cells/mL, approximately 5-fold higher than that of normal peripheral blood. The RBC content of leukopaks was tightly maintained, with a hematocrit below 2% (mean hematocrit: 1.9 ± 0.3%) (**Fig. 2C-H**).

**Figure 1:**
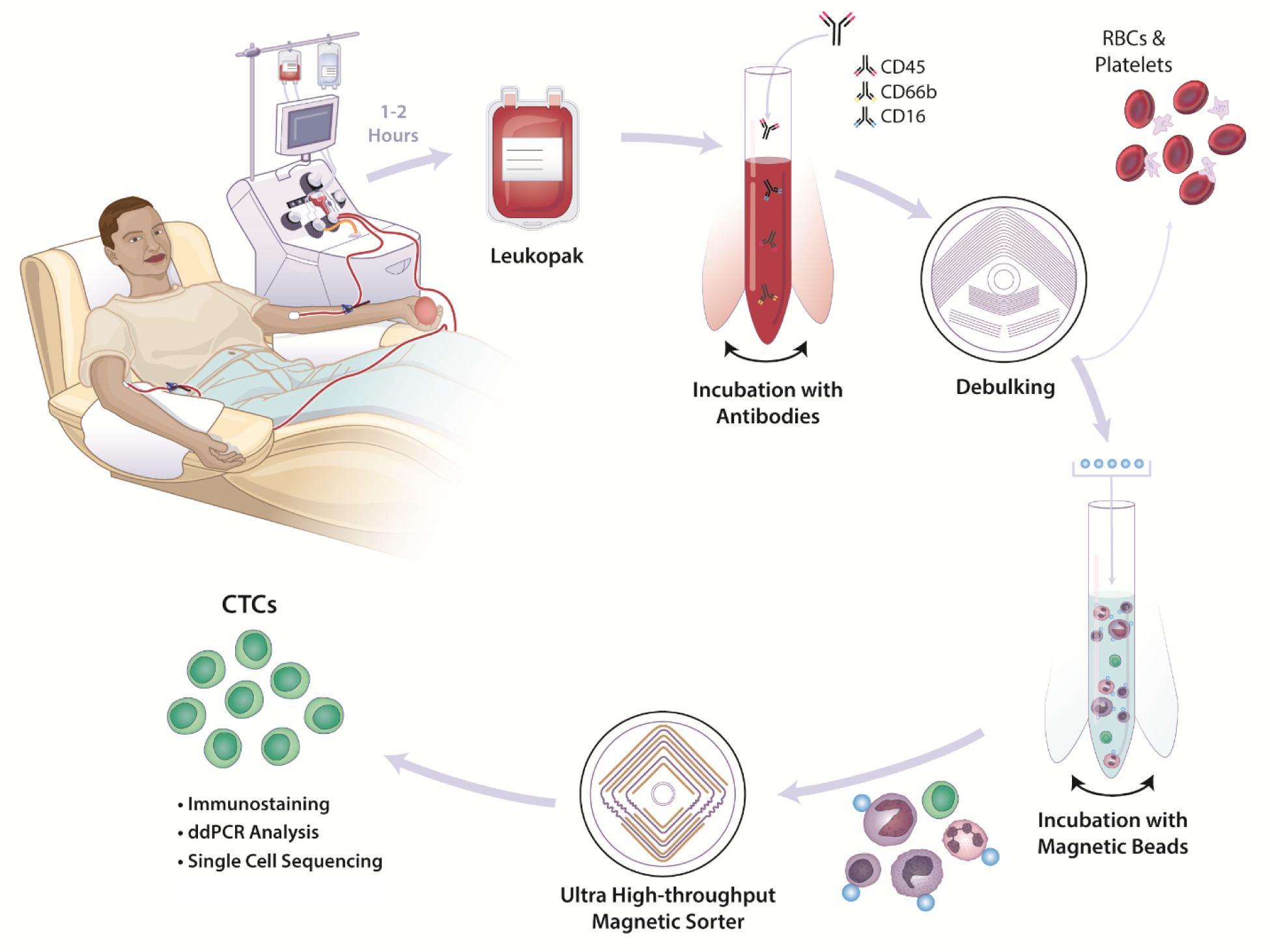
Schematic illustration of high-volume CTC enrichment using microfluidic isolation of leukapheresis samples. The leukapheresis product (leukopak) is collected from a patient during a 1 to 2 hour procedure that samples 3 to 6 L of blood volume. Following the addition of biotinylated antibodies to tag WBCs, a microfluidic debulking chip is used to remove unbound antibodies, RBCs, platelets, and excess plasma. Streptavidin-conjugated magnetic beads are added to tag WBCs, which are then separated from untagged CTCs using an ultrahigh-throughput microfluidic magnetic sorter. Enriched CTCs are imaged using immunofluorescent staining for lineage or tumor markers, subjected to RNA-based quantitation of specific transcripts (ddPCR), or cultured ex vivo. Single CTCs may be isolated for RNAseq or DNA analyses, including mutational profiling and CNV analyses. Together, the two chips comprise the ^LP^CTC-iChip platform.

**Figure 2:**
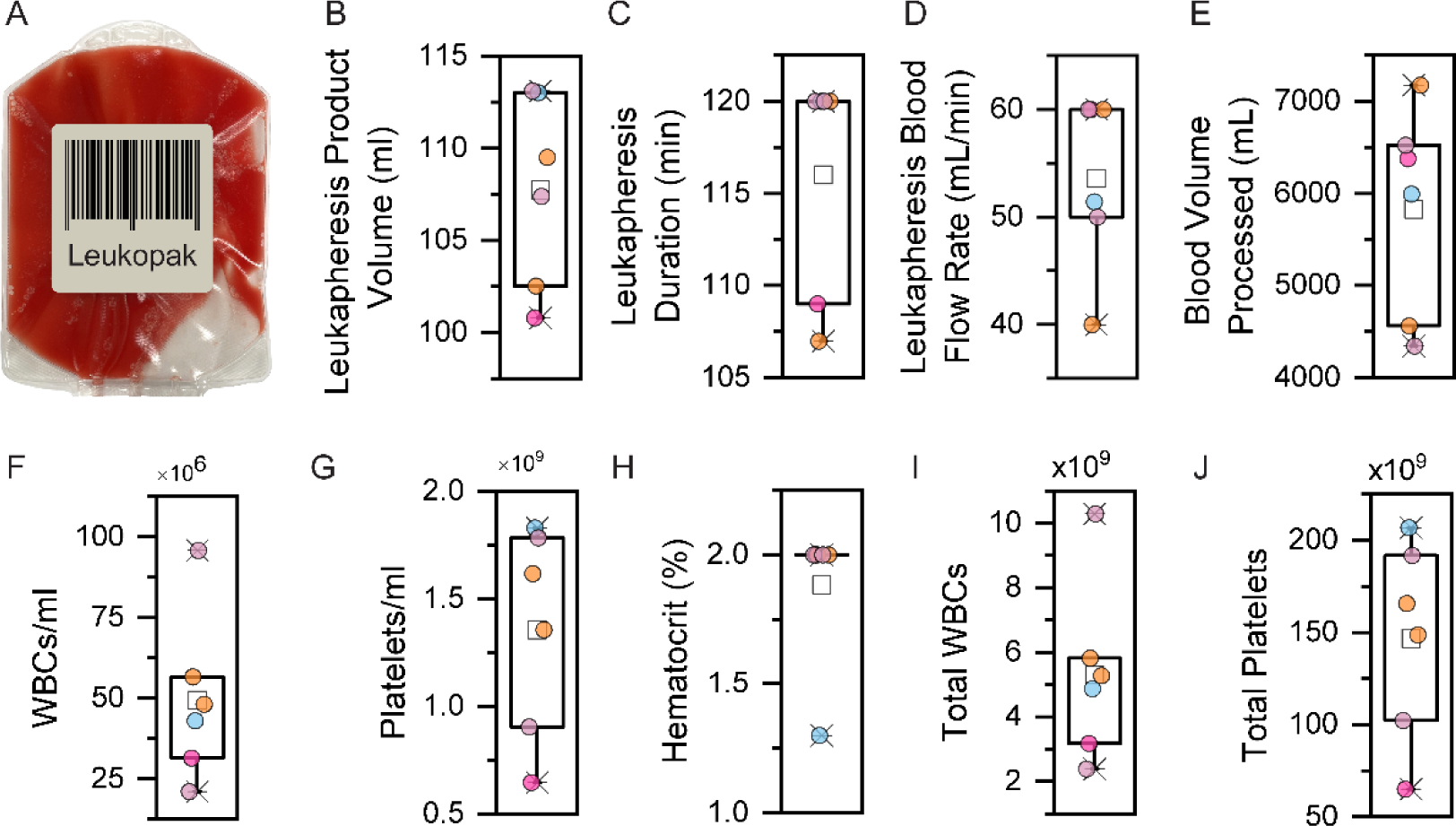
Patient leukapheresis parameters and cellular content in the six cancer patient-derived leukopaks. (A) Representative image of a 100 mL leukopak obtained from a 2-hour leukapheresis session. (B-E): Leukapheresis product volume (B), procedure duration (C), blood flow rate (D), and processed blood volume through the apheresis machine, respectively (E). (F-H): Concentration of WBCs (F), platelets (G) and hematocrit (RBCs; H) in leukopak samples. (I-J) Total number of WBCs (I) and platelets (J) in cancer patient-derived leukopaks, containing a median 88-fold more WBCs and 49-fold more platelets, compared with a standard 10 mL whole blood sample.

In addition to the high concentration of WBCs and platelets within 100 mL leukopaks, their total numbers were considerable, with a mean of 5.3 ± 2.5 billion WBCs and 146.7 ± 49.4 billion platelets, representing 88-fold and 49-fold, respectively, larger amounts than present in a typical 10 mL blood sample used for microfluidic CTC enrichment (**Fig. 2I-J**).

### Microfluidic CTC enrichment from patient-derived leukopaks

In testing large blood volumes spiked with cultured CTCs, we previously applied two microfluidic devices in series: an initial debulking chip to remove RBCs and platelets (inertial separation array chip), followed by a magnetic lens-based high-throughput cell sorter (MAGLENS) (**Fig. 3 and Sup Fig 1**)^22,23^. We used the same microfluidic devices to process patient-derived leukopaks, designed to operate in the “negative depletion mode,” removing massive numbers of normal blood cells to purify untagged and unmanipulated CTCs. To achieve this, we first briefly incubated the entire leukopak specimen with biotinylated antibodies directed against the common leukocyte markers CD45 (pan-leukocytes), CD66b (granulocytes), and CD16 (monocytes), and then flowed the leukopak through a microfluidic debulking chip to remove excess free antibodies, along with plasma, RBCs, and platelets (**Fig. 1**).

**Figure 3:**
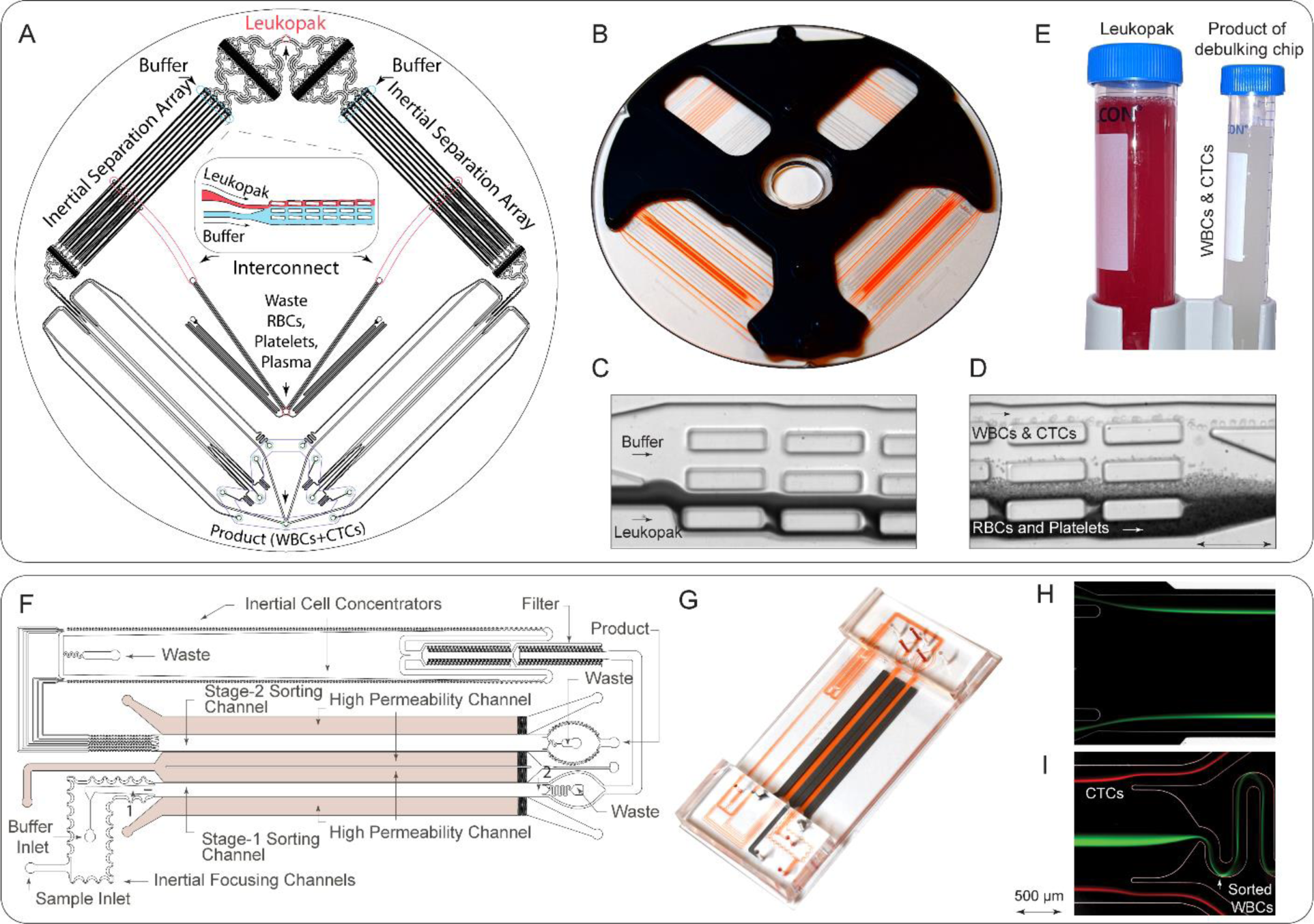
High-throughput microfluidic devices used for processing leukopaks. (A, B) Schematic (A) and image (B) of the debulking chip used for removing unbound antibodies, RBCs, platelets, and excess plasma, concentrating WBCs and CTCs into a clean buffer. (C, D) Images illustrating the inertial separation of nucleated cells from RBCs and platelets within the debulking chip. The underlying inertial separation array technology^23^ repeatedly deflects larger nucleated cells over an array of rectangular islands (microposts) using wall lift forces to transfer the cells into a clean buffer stream, separating them from the smaller cells and non-nucleated cells (RBCs, platelets). Following the entry into the device (C), serial deflection across the multiple islands that comprise the Chip allows the collection of nucleated cells with very high purity and yield (D). (E) Image of the input sample (leukopak) before debulking, with red color illustrating high RBC content, and of the purified product after debulking of RBCs and platelets. (F) Design of the microfluidic magnetic sorter for depleting magnetically labeled WBCs, using cascaded two-staged magnetic sorting and a very high-gradient magnetic field, which is created by magnetic lenses adjacent to the cell flow channels^22^. (G) Image of the microfluidic magnetic sorter device. (H, I) Streak images of cells at the inlet (H) and exit (I) of stage I of the magnetic sorter. Magnetic bead-labeled white blood cells (green) are deflected into the central core of the sorting channels, away from CTCs (red), thereby allowing continuous depletion of WBCs without clogging the channel.

The debulking chip takes advantage of inertial flow-based size separation using an array of rectangular microposts (200µm × 50µm × 52µm), as previously described (**Fig. 3A-E**) ^22,23^. Using co-flow principles, the stream of fluid from the leukopak specimen is directed close to the wall of the microposts (**Fig. 3A**), where WBCs and CTCs experience a higher wall lift force due to their larger size, thereby moving further away from the wall compared with RBCs and platelets (**Fig. 3C-D**). The stream of fluid (3.6%) near each rectangular wall of microposts that contains RBCs and platelets is then siphoned away, a process that is repeated over 180 serial microposts across the entire array, thereby enabling highly efficient removal of RBCs (99.95%) and platelets (99.98%) while achieving a high yield of WBCs and CTCs, consistent with our previous studies (**Fig. 3E**)^22^. To achieve efficiency, we parallelized 16 such inertial separation devices in a plastic disk that we call the ^LP^debulking chip, as shown in **Fig. 3A-B**. We used 3 to 4 ^LP^debulking chips to process a single leukopak, depending on the total volume. This optimization achieved a leukopak sample flow rate ranging from 109.5 mL/h to 146 mL/h and a total buffer flow rate from 522 mL/h to 696 mL/h.

Following leukopak debulking, we incubated the WBC and CTC suspension with 1 µm streptavidin-conjugated magnetic beads before flowing these through two parallel microfluidic MAGLENS sorters to deplete bead-bound WBCs (**Fig. 3F-G**). The high concentration and large total number of WBCs in leukopaks require a 50-times higher cell processing throughput than the existing state-of-the-art magnetic sorters^7,20,24,25^. Since the flow throughput of a continuous magnetic flow sorter is directly dependent on the magnetic field gradient in the deflection channels, effective processing relies on the “magnetic lenses” within the MAGLENS sorter design. As previously described^22^, these are created by the incorporation of high-permeability channels filled with soft magnetic iron particles, that amplify the magnetic field gradient by 35-fold. The micromagnetic lenses, which are lithographically defined within 60 μm of the cell sorting channels, create a field gradient as high as 15,400 T/m compared to 440 T/m created for the conventional CTC-iChip arrangement used to process a standard 10 mL tube of whole blood^7,20,22^ (**Sup Fig. S2**). Utilizing these principles, each MAGLENS sorter can process a leukopak sample at a flow throughput of 48 mL/h (3 billion cells/h), 60-times higher than the conventional CTC-iChip. Given the very high number of WBCs to be depleted from leukopaks and the range of expression of leukocyte markers, we applied a cascaded two-staged magnetic sorting system. In stage I, cells tagged with >10 beads are deflected, and in stage II, WBCs labeled with a single magnetic bead are removed^22^ (**Fig. 3F-G**). For the initial stage I depletion of highly tagged cells, the two asymmetric serpentine channels apply inertial focusing to generate a single-cell stream by balancing shear-induced lift and Dean flow-based drag forces (**Fig. 3H**). This inertial focusing arranges single cells in a line, minimizing the possibility of many WBCs colliding with a CTC during sorting while magnetic forces deflect tagged cells toward the center streamline of the channel (waste). **Fig. 3I** shows streak images of green-fluorescent WBCs efficiently sorted from red-fluorescent CTCs. Effective depletion of WBCs carrying fewer magnetic beads requires stage II, where the flow rate is reduced through on-chip concentration to enable the depletion of cells labeled with a single bead. This is accomplished through an inertial cell concentrator^22^ before stage II of the magnetic sorter, thereby achieving an 11-fold concentration of the final product. Both the co-flow in stage I and the inertial concentrators in stage II ensure that cells remain close to the side walls, where the magnetic force is strongest due to proximity to magnetic lenses (**Sup Fig. S2**). Overall, the total flow rate across Stages I and II of the MAGLENS chip, including both leukopak sample and the added buffer, is 168 mL/h.

Magnetic forces in both stages of the sorter are designed to deflect labeled cells into the center of the channel, where magnetic forces vanish (**Fig. 3I**). This unique design precludes any possibility of clogging as cells are moved into the core of the flow away from walls. Extending from our initial prototype^22^, we made two significant adjustments to the magnetic sorter to successfully process patient-derived leukopaks. Instead of using an on-chip stage I filter, we designed a completely new filter chip with an aperture size of 42 µm to allow filtration of any large aggregates of leukocytes or clots that may form in patient-derived leukopaks (**Sup Fig. S3**). We also added DNAse at 100 units/mL to the cell suspension to digest any neutrophil extracellular traps (NETs), which cause fouling of the microfluidic features. These strategies now allow us to achieve clog-free sorting while handling billions of nucleated cells. Having demonstrated the efficacy of the two optimized components of the ^LP^CTC-iChip (Debulking and MAGLENS chips), the system is now capable of being integrated into a single device for future point-of-care applications.

### Multispectral imaging analysis of patient leukopak-derived CTCs

To identify CTCs within the final microfluidic products, we first applied immunofluorescence (IF)-based staining and high-content multispectral imaging, followed by digital image processing and manual validation. CTCs were defined as being positive for both nuclear signal (DAPI) and the relevant characteristic tumor markers (see below), but negative for the cocktail of WBC markers (CD45, CD66b, and CD16). Antibodies against tumor markers used for the different tumor histologies were a cocktail of: EpCAM, pan-cytokeratin (pan CK) and CK19, for prostate cancer (GU) and triple negative breast cancer (TNBC)^7^; EpCAM, pan CK, CK19, asialoglycoprotein receptor 1 (ASGR1) and Glypican 3 (GPC3), for hepatocellular carcinoma (HCC)^26^; and Sox10, melanocyte differentiation antigen (Melan-A), and neuron glial antigen-2 (NG2), for uveal melanoma (UM)^27^. Antibodies against the three WBC antigens were grouped in one fluor (Alexa Fluor 647), and those against the various tumor antigens in a second fluor (Alexa Fluor 488) (**Fig. 4A-E**). Using these criteria, we calculated a median yield per leukopak of 2,799 CTCs (GU-1: 58,125 CTCs, TNBC-1: 4,580 CTCs, HCC-1: 4,109 CTCs, HCC-2: 1,490 CTCs, UM-1: 720 CTCs, UM-2: 100 CTCs (**Fig. 4F**). The average WBC depletion was 99.96% (3.4 ± 0.3 Log_10_ depletion), thereby achieving a CTC purity ranging from 0.005% (UM-1) to 3.3% (GU-1) (**Sup. Fig. S4A and B**).

**Figure 4:**
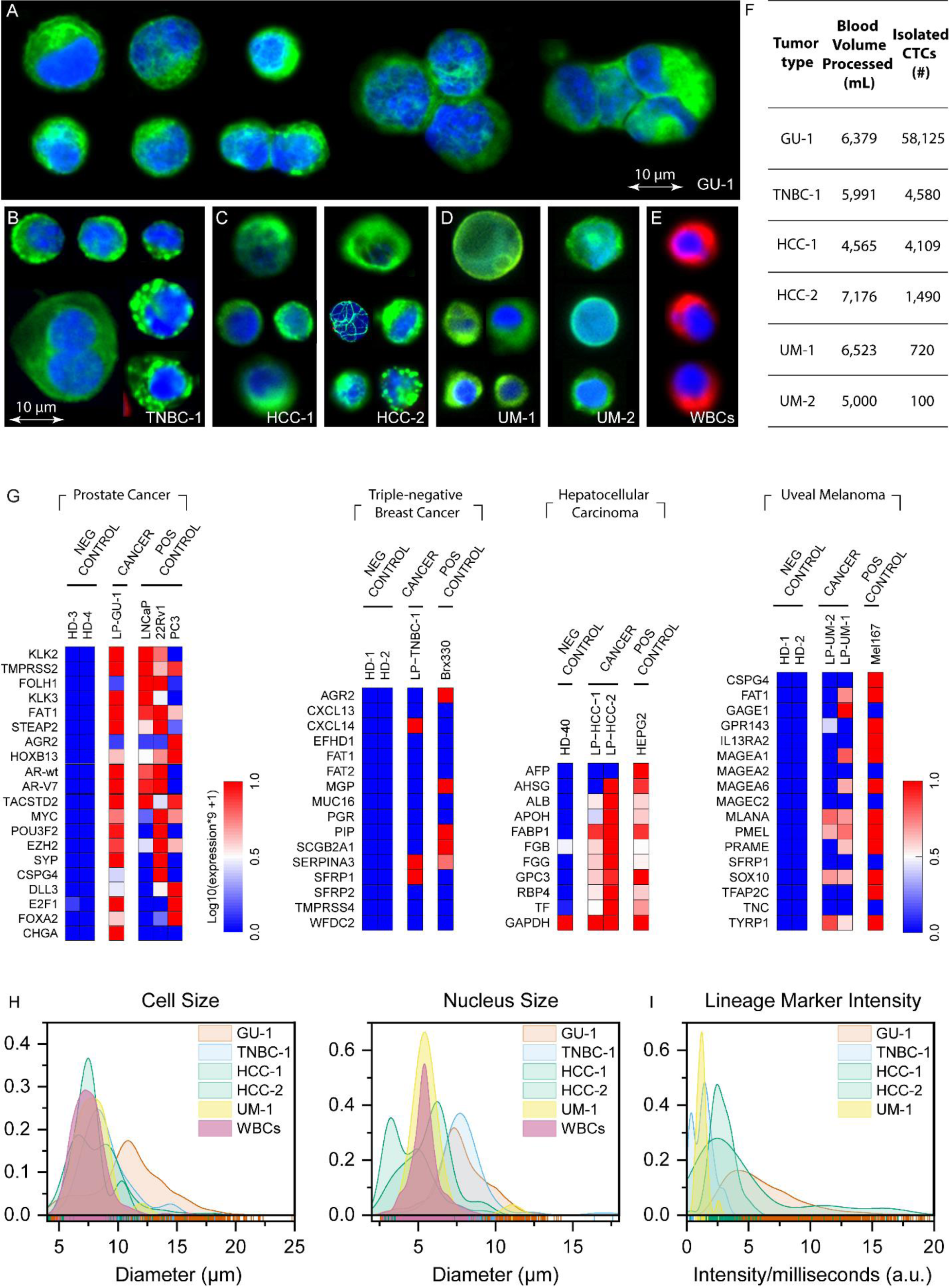
Analyses of microfluidic-enriched CTC bulk populations. (A-D) Immunofluorescence images of representative CTCs enriched from leukopak samples from patients with metastatic prostate cancer (GU-1), triple-negative breast cancer (TNBC-1), hepatocellular carcinoma (HCC-1 and HCC-2), and uveal melanoma (UM-1 and UM-2). Bulk cell populations are stained with DAPI nuclear marker (blue), the relevant tumor markers grouped within a single color (green) (see individual epitopes below), and with antibodies against the WBC markers CD45, CD16, CD66b (red). Tumor markers: GU-1 (A; EpCAM, pan CK, CK19), TNBC-1 (B; EpCAM, pan CK, CK19), HCC-1 and HCC-2 (C; EpCAM, pan CK, CK19, ASGR1, GPC3), UM-1 and UM-2 (D; Sox10, Melan-A, NG2)). Representative contaminating WBCs are shown in (E). (F) Table listing blood volumes processed, and yield of CTCs obtained from analysis of entire patient-derived leukopaks (median 2,799 CTCs per leukopak). (G) Droplet digital RNA-PCR (ddPCR) analysis of 0.5 to 1% of the bulk CTC products from all six cases, quantifying expression of previously curated RNA signatures that denote either tissue lineage or cancer-specific transcripts within the background of normal blood cells. Of note, GU-1 CTCs express multiple neuroendocrine genes (CHGA, SYP, DLL3), consistent with immunohistochemistry staining for synaptophysin and chromogranin A in a resected adrenal metastasis from this patient. Healthy donor blood is shown as negative control, with positive controls drawn from either cultured prostate cancer cell lines (LNCaP, 22Rv1, and PC3), cultured breast CTCs (BRx-142^33^), cultured liver cancer cells (HepG2), or cultured melanoma CTCs (Mel-167^48^), respectively. (H) Measured whole cell and nucleus diameters of individual CTCs, compared with WBCs. A total of 5,543 CTCs were visualized (GU-1, TNBC-1, HCC-1, HCC-2, UM-1) and compared with 223 WBCs from the five cases, all of which were present in the final microfluidic product (selected using negative depletion of most WBCs and hence independent of CTC size bias). Substantial overlap in size is evident between CTC and WBC populations. Considerable heterogeneity in both cell and nucleus size is also present within the CTCs of individual patients, as is a bimodal distribution in cell and nucleus size in the CTCs of both HCC patients. (I) Variation across individual CTCs from cases GU-1, TNBC-1, HCC-1, HCC-2, and UM-1 in their intensity of staining for the combined lineage markers (see above), as imaged using Vectra Polaris multispectral imaging system and HALO quantitative image analysis platform.

To confirm the tissue of origin of the enriched CTCs, we removed small aliquots, ranging from 0.5 to 1% of the final microfluidic products, to apply previously defined digital, cell linage-based RNA signatures that capture the heterogeneity of prostate cancer (20 genes), breast cancer (17 genes), liver cancer (11 genes) and melanoma (19 genes) CTCs, while distinguishing them from surrounding blood cells with very high specificity^27–30^. In all six cases, droplet digital PCR (ddPCR) assays confirmed the presence of the expected tumor type (see **Fig. 4G**). As negative controls, healthy donor blood processed through the same microfluidic platform showed no ddPCR signal; diluted RNA from histologically relevant cancer cell lines was used as positive control.

Studies of heterogeneity across CTCs have been hampered by the small number of cancer cells isolated from individual patients using current technologies, thereby confounding inter-and intra-patient variability. The ability to isolate such large numbers of CTCs from individual patient-derived leukopaks can thus provide a vastly improved initial measure of true intra-patient CTC diversity. Enrichment of CTCs through negative depletion of WBCs is relatively unbiased compared with positive selection of CTCs based on their expression of pre-selected epithelial markers such as EpCAM or the application of size-based selection that assumes epithelial cells to be larger than leukocytes. Indeed, morphological analysis of all scored CTCs from the patient samples, compared with their matched WBCs, shows substantial overlap in their cell diameter, with 99% of WBCs similar in size to 66% of CTCs (**Fig. 4H**). These findings are consistent with the relatively poor yield of size-based CTC isolation across different cancer histologies. Within the many CTCs isolated from individual patients, we also note considerable variation in nuclear and cell diameters, with CTC diameters ranging from 4 µm to 25 µm (**Fig. 4H**). Interestingly, a bimodal distribution in both nuclear and total cell size is evident in the two HCC cases (**Sup. Fig. S4C).**

We also used multispectral imaging to quantify the variability in tumor marker expression across the large number of CTCs isolated from individual patients (**Fig. 4I and Sup. Fig. S4D**). The cocktail of tumor markers used to identify CTCs within each tumor type provides an initial measure of both lineage and epithelial differentiation. The variation in tumor marker expression ranged from 3.7-fold in UM-1 to 18.8-fold in HCC-2. Similar analyses may be used to ascertain the heterogeneous expression of selected antibody-drug conjugate (ADC) markers on CTCs within individual patients in the context of personalized antibody-directed cancer therapies.

### Single-cell analysis of CTCs for whole genome and transcriptome sequencing

On-treatment tumor biopsies have provided invaluable information about the mechanisms whereby cancers acquire drug resistance, including novel mutations, altered signaling pathways, and even cell lineage alterations. Blood-based ctDNA sequencing may reveal acquired mutations within selected cancer gene panels but given low quantities of small molecular weight DNA at low purity, it cannot achieve sufficient coverage for whole exome sequencing, nor can it provide RNA sequencing to identify altered transcriptional programs. The large number of intact CTCs isolated from leukopaks thus offers a unique opportunity to combine non-invasive blood-based diagnostics with detailed interrogation of cancer cells at the single-cell level.

As proof-of-principle, we selected the prostate cancer case GU-1, in which a large number of CTCs (58,125 CTCs) made it possible to optimize molecular analyses. To readily isolate single CTCs from the microfluidic ^LP^CTC-iChip enriched population (3.3% purity), we processed cells through a second purification step, applying a microfluidic fluorescence-activated cell sorter (SONY SH800) to distribute individual CTCs into single wells. We used AF488-conjugated antibodies against both the epithelial marker EpCAM and the prostate-specific markers PSMA *versus* PE-Cy7-conjugated antibodies targeting the WBC markers CD45, CD16, and CD66b to distinguish prostate CTCs from contaminating blood cells, enabling efficient high throughput single-cell sorting (**Fig. 5A, Sup. Fig. S6A**). To further optimize the sorting process, we implemented a sequential sorting method, with an initial high-yield bulk sorting step of AF488-tagged CTCs and removal of dead cells and cell fragments, followed by sorting single cells into individual wells of PCR plates (**Fig. 5B**). Within the individual wells, we separated intact nuclei from single cells for DNA sequencing and the corresponding cytoplasm for RNAseq analysis, generating templates for paired single-cell whole genome sequencing (WGS) and single-cell RNA sequencing (RNA-seq, Smart-seq2 method)^31^.

**Figure 5:**
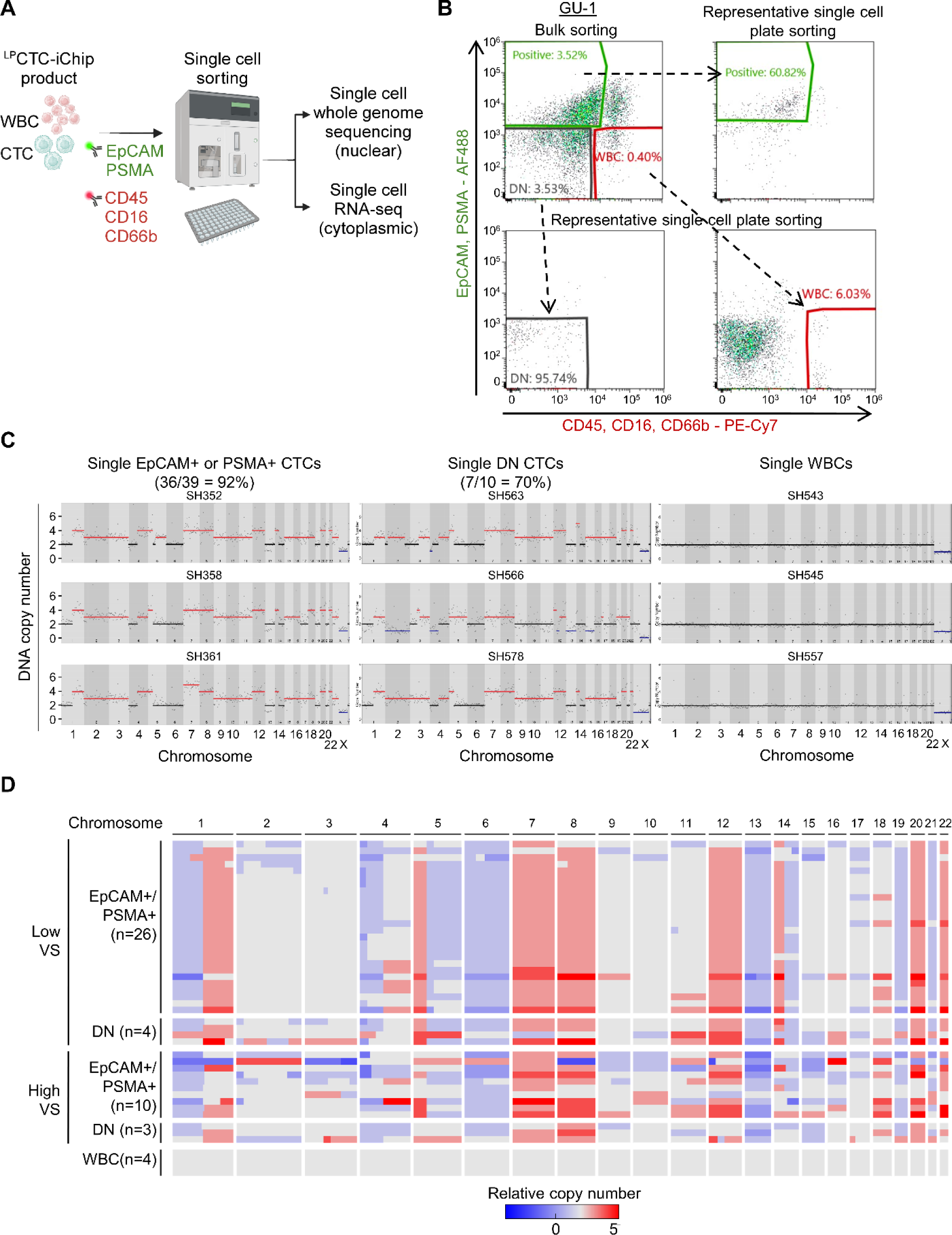
Single-cell whole genome sequencing and DNA copy number analyses. (A) Schematic of single-cell isolation from ^LP^CTC-iChip-enriched leukopak samples using Fluorescence Activated Cell Sorting (FACS). The CTC-enriched ^LP^CTC-iChip product is free of magnetic-conjugated antibodies against WBC markers CD45, CD16 and CD66b. For FACS single-cell isolation, two-color separation is achieved using an AF488-conjugated antibody cocktail against EpCAM and PSMA (for GU-1; green) and a PE-Cy7-conjugated antibody cocktail against CD45, CD16, CD66b (labeling contaminating low-expressing WBCs that escaped ^LP^CTC-iChip depletion; red). Individual cells (CTCs and WBCs) are then subjected to paired single-cell whole genome sequencing and single-cell RNA-seq. (B) Two-step FACS-sorting strategy using an initial ultra-yield bulk sorting to remove dead cells and debris, followed by single-cell plate sorting (representative figure of single cell sorting into a single 96-well PCR plate shown here), to isolate candidate CTCs that are positive cells for either EpCAM (epithelial marker) or PSMA (prostate lineage marker); cells of uncertain identity that are negative for both EpCAM and PSMA, as well as the WBC markers CD45, CD16 and CD66b (Double Negative; DN); and white blood cells that are positive for CD45, CD16 and CD66b. 576 candidate CTCs and 192 DN cells were collected using high-purity single-cell sorting. (C) Representative DNA copy-number variation (CNV) analysis in individual CTCs, compared with diploid WBCs from the same patient (X chromosome haploid in male patient). Cancer cell-associated aneuploidy was confirmed by CNV in 36/39 single CTC candidates (EpCAM or PSMA positive) and in 7/10 cells of uncertain identity (Double Negative). Ginkgo was used for DNA copy number analysis from single-cell whole genome sequencing data. (D) Normalized heatmap showing clustering of single-cell CNVs across the genome in CTCs (n=43) from patient GU-1. All CTCs show shared core chromosomal alterations, WBCs are diploid. Ginkgo was used for DNA copy number analysis. Variability score (VS) is used to quantify DNA and assay quality (low VS corresponds to high quality).

We sorted 576 individual candidate CTCs using high-purity single-cell sorting, based on their expression of the epithelial marker EpCAM or the prostate lineage marker PSMA (combined AF488 staining), and absent expression of multiple WBC markers (combined PE-Cy7 staining). In addition, we collected 192 “double negative” cells, lacking staining for either AF488 or PE-Cy7). Such double negative (DN) cells are abundant in CTC-enriched populations derived from the depletion of WBC markers, but their identity is uncertain. We selected 84 single cells (57 AF488-positive, 27 DN) for Whole Genome Sequencing (WGS) using the MALBAC method^32^, with 1.5-2x genome coverage depth (Ginkgo tool, http://qb.cshl.edu/ginkgo), allowing for analysis of DNA copy number variation (CNV), a definitive marker of cancer-related aneuploidy. As controls, single CTCs from *ex vivo* cultured breast CTCs lines^33^ showed characteristic CNV, while single WBCs contaminating the microfluidic preparation showed the expected normal diploid genomes (**Sup. Fig. S6C, D**). After filtering out samples with low-quality DNA sequencing, CNV analysis indicated high confidence aneuploidy in 36/39 (92%) single cells expressing EpCAM or PSMA and in 7/10 (70%) double negative cells. These alterations included shared genomic amplifications and deletions across most CTCs, as well as some unique chromosome changes consistent with tumor cell heterogeneity (**Fig. 5C, D**). The CNV pattern of gains and losses among EpCAM/PSMA positive cells and the double negative cells was indistinguishable, indicating that these DN cells are genuine CTCs that have lost epithelial and prostate lineage markers. Key chromosomal alterations universally detected in CTCs included copy number gains in chromosomes 1q, 7, and 8, as well as losses in chromosomes 1p, 5, 6, and 13, which are commonly observed in advanced prostate cancer, as reported in the TCGA and SU2C cohorts^34–36^ (**Fig. 5D**).

While the high number of CTCs recovered from GU-1 made it possible to perform detailed CNV analyses, we were also able to demonstrate aneuploidy within single circulating cancer cells in the two HCC cases which had fewer CTCs (**Sup. Fig. 7A, B**). Whereas GU-1 CNV analysis indicated a single dominant genomic lineage, the 12 single CTCs analyzed from HCC-2 showed two distinct subpopulations (n=7 vs. 5), with a subset of shared core genomic amplifications and deletions, as well as clustered unique chromosome changes consistent with genomic heterogeneity in metastatic cancer (**Sup. Fig. 7C, D**). The shared chromosomal alterations identified include copy number gains in chromosomes 1, 6, and 8 and copy number losses in chromosomes 4, 13, 16, and 17, which are prevalent in hepatocellular carcinoma^37–39^ (**Sup. Fig. 7D).**

In case GU-1, we applied paired single cell RNA-seq and CNV analyses within individual CTCs, separating cytoplasm for RNA analysis from nucleus for DNA analysis (**Fig. 5A, Sup. Fig. S6C, D**). Among CTCs with definitive CNV, 30 cells retained sufficiently high-quality RNA for single-cell RNA-seq. In these cells with a shared, clonal CNV pattern, unsupervised clustering analysis revealed two distinct hierarchical clusters based on RNA expression (**Fig. 6A**). Unsupervised gene set enrichment analysis (GSEA) identified a striking upregulation of FGFR signaling in Cluster-1, while Cluster-2 showed upregulation of pathways associated with inflammatory responses, chemokine signaling, IL/JAK/STAT signaling, cell adhesion, and epithelial-mesenchymal transition (FDR ≤ 0.25) (**Fig. 6B, Sup. Table S4**). In addition, compared with Cluster-2, Cluster-1 demonstrated enrichment in prostate cancer-associated pathways, including androgen receptor (AR) signaling, proliferation, and DNA repair pathway (**Fig. 6C**).

**Figure 6:**
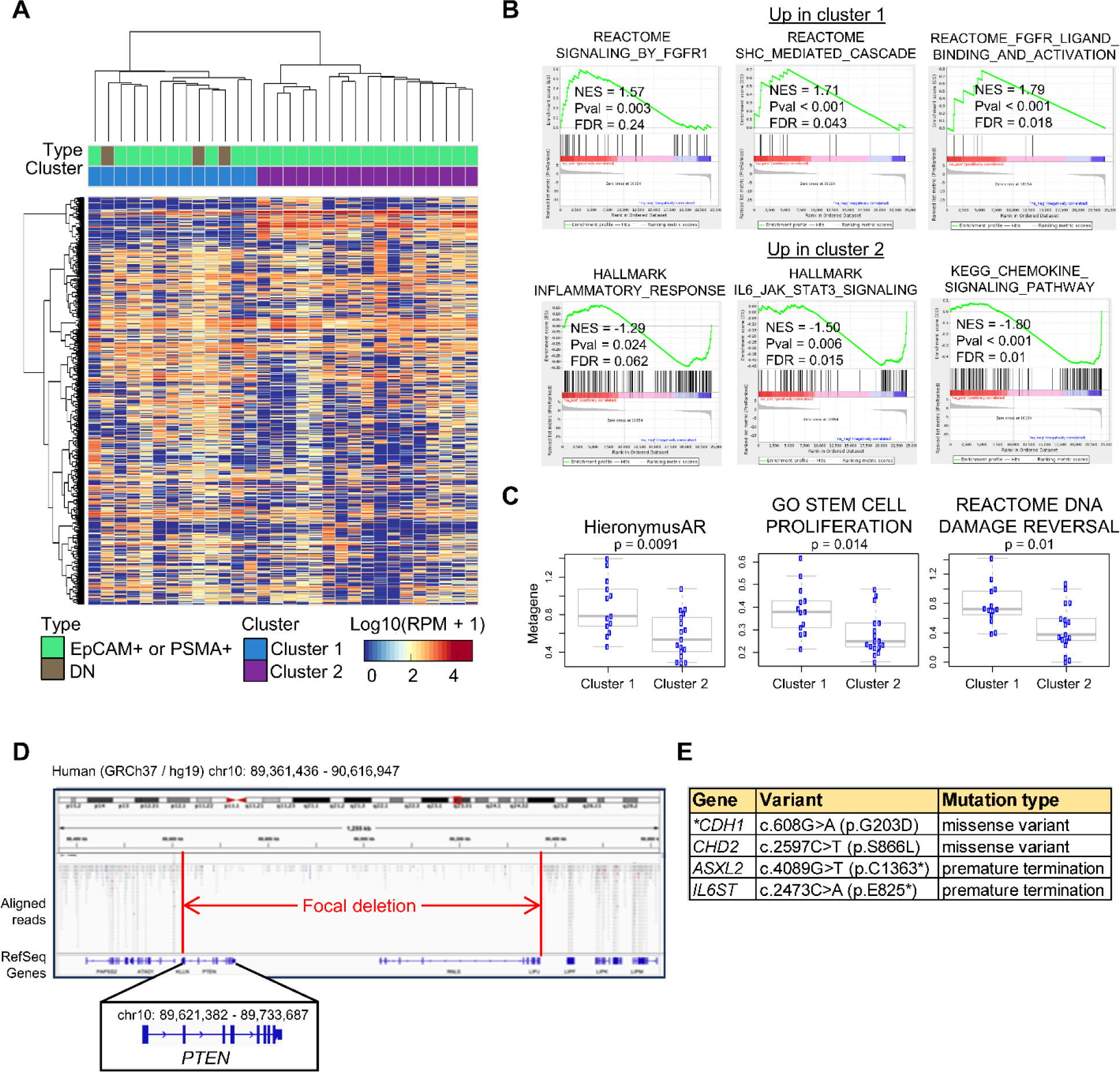
Single-cell RNA-seq and whole exome sequencing (WES) data analysis for GU-1 prostate CTCs. (A) Heatmap showing unsupervised hierarchical clustering of single-cell RNA-seq data, showing two distinct expression clusters. The three Double Negative (DN) cells fall within cluster-1. (B) Unsupervised gene set enrichment analysis (GSEA) shows elevated FGFR signaling in cluster-1 (three different signatures), whereas cluster-2 shows enrichment in inflammatory signaling pathways. Permutation-based P-value determined by a normalized enrichment score (NES) and FDR value from GSEA software. (C) Supervised GSEA, showing enrichment in cluster-1 for androgen-induced and repressed genes (Hieronymus AR), cell proliferation, and DNA repair pathways (D) IGV visualization of a deletion spanning the PTEN gene, identified by CTC WES sequencing. The PTEN deletion was also detected in a matched surgical resection of a metastatic tumor lesion. (E) Presumed clinically significant mutations identified by CTC WES sequencing and analyzed using MuTect software. The CDH1 mutation (asterisk) was also reported in the matched metastatic lesion (cancer gene-specific panel), but the other three mutations were uniquely identified by WES analysis of CTCs.

To extend from CNV analysis of GU-1 CTCs to sequencing for point mutations, we undertook whole exome sequencing (WES). Since single-cell WES may not provide adequate coverage for comprehensive mutational analysis, we pooled 26 CTCs displaying CNV (AF488-positive) and subjected them to “pseudo-bulk” DNA sequencing. WES analysis of CTCs revealed a focal homozygous deletion spanning the *PTEN* gene and a *CDH1* G203D mutation, both of which were the only genetic alterations that had been identified in an adrenal metastasis previously excised from the patient and analyzed using the FDA-approved FoundationOne®CDx test. In addition, CTC WES analysis revealed a likely functionally significant missense mutation in *CHD2*, and stop codons in *ASLX2* and *IL6ST*, along with variants of uncertain significance within known cancer genes (**Fig. 6D, E, Sup. Table S5**).

Taken together, the combination of high throughput microfluidic processing of leukopaks for large-scale CTC enrichment with high-efficiency single-cell sorting for quantification of CNV, point mutations, and gene expression enables serial non-invasive monitoring of multiple parameters that may help guide therapeutic decisions.

## Discussion

We have described the clinical application of a novel high throughput microfluidic device for efficient and semi-automated enrichment of large numbers of CTCs from leukopaks drawn from patients with metastatic cancer. To date, CTC analyses have had limited clinical utility, given the extremely rare number of cancer cells in the blood circulation of patients with cancer and the fact that current technologies can only process 10 to 20 mL of blood for CTC enrichment. As such, the ability to consistently obtain large numbers of cancer cells through microfluidic enrichment of an entire leukapheresis product presents a unique opportunity to transform both research and clinical applications of CTCs in cancer.

### Microfluidic processing of large cell numbers and volumes

Leukapheresis is routinely used in clinical settings to obtain sufficient numbers of hematopoietic stem cells for bone marrow transplantation or T cells for CAR-T engineering^14,19^. In these applications, normal subsets of hematopoietic mononuclear cells are enriched through centrifugation via leukapheresis, followed by immunological selection, achieving purities of >90% and without requiring downstream single-cell analytics. In contrast to these hematopoietic cells, CTCs are considerably rarer in the bloodstream and are often pre-apoptotic from loss of cell adhesion signals, shear stresses, and oxygen tension within the vasculature. The robust isolation of large numbers of CTCs from patient-derived leukopaks thus presents a major technological challenge, both in terms of ultra-rare cell detection as well as the need for low-shear stress microfluidics. Furthermore, the most insightful results to be derived from CTC analyses stem from single-cell-level molecular analyses, in which tumor heterogeneity at DNA, RNA, and protein levels can be assessed.

To our knowledge, real-time flow microfluidic devices have never been applied to processing the very large and concentrated cell volumes described here. From a technological standpoint, microfluidic processing of leukopaks for ultra-rare cell detection is hampered by the large total number of WBCs (approximately 6 billion cells), their high concentration (8-fold higher than normal blood), and the large total fluid volumes (100 mL). Clogging of channels, resulting from platelet aggregation or release of DNA NETs from lysing WBCs, is a major obstacle (**Sup. Fig. S5**), as is target cell loss from the massive numbers of non-target cells to be removed and associated prolonged processing time. Applications of current CTC enrichment technologies to analyze small fractions of leukopaks^15,16,40^ have confirmed predictions that processing the entire leukapheresis product would indeed generate very large numbers of CTCs, but such complete leukopak sorting technologies have not been described. Compared with batch purification strategies, continuous flow processes through microfluidic platforms have the advantage of gentle processing, and the inertial focusing platform that we have developed is capable of very high CTC capture rates since individual cells flow in a single file as they are interrogated through a high-gradient magnetic deflection field.

Two essential features in the ^LP^CTC-iChip make efficient sorting of large cell volumes possible. First, both components of the chip are designed to deflect blood cells away from the walls, thereby minimizing the possibility of clogging despite the very high concentration of WBCs and platelets. In the debulking chip, the cell-free region near the walls of rectangular pillars is created by wall lift forces (**Fig. 3C-D**). In the MAGLENS sorter, the magnetic field directs magnetically labeled cells into the core of flow at the center of the channel, away from the side walls (**Fig. 3I**). This is made possible by the coplanarity of the opposing magnetic lenses on either side of the sorting channels, with a symmetric force towards the center of the channel, such that the magnetic gradient gradually vanishes as cells near the center of the channel (**See Sup. Fig. 2**). As a consequence, the sorted cells are moved into the core of the Poiseuille flow, creating an inherently clog-free, safe design that can process billions of cells and a large fluid volume.

Second, the MAGLENS sorter employs a two-stage sequential strategy to deplete WBCs, a critical consideration since WBCs are highly variable in their expression of the cell surface markers used to bind magnetic-conjugated beads. As such, WBCs with high expression of CD45, CD16, or CD66b acquire large loads of magnetic beads on their cell surface during labeling, and these cells may clog channels upon application of a strong magnetic force. The Chip design thus involves an initial deflection of WBCs with a large magnetic load through a high-flow channel in Stage I, followed by the passage of remaining WBCs with lower epitope expression and magnetic load through a low-flow channel in Stage II. Approximately 90% of WBCs are removed in stage I, with the rest in stage II, achieving a 99.96% total depletion without compromising the flow rate or risk of clogging.

We note that the ^LP^CTC-iChip platform enriches CTCs through “negative depletion” of WBCs and is therefore tumor epitope agnostic. This is illustrated in the tumor types selected for analysis here: melanomas, including uveal melanomas (UM), have a minimal expression of traditional epithelial markers such as EpCAM, and both hepatocellular cancers (HCC) and triple-negative breast cancer (TNBC) also have relatively low EpCAM expression. For these tumor types, the initial epitope-agnostic enrichment is followed by CTC scoring by staining with a cocktail of antibodies targeting multiple shared and lineage-specific epitopes. Nonetheless, the ^LP^CTC-iChip platform used here is also adaptable to a “positive selection” mode, including the capture of CTCs by virtue of their expression of EpCAM or other epithelial or tumor-specific epitopes, which may be valuable in specific tumor-targeting contexts.

While discrimination between rare CTCs and abundant surrounding WBCs using antibody-mediated detection has limitations, including variability in cell surface protein expression, it appears well suited for large-volume enrichment strategies. Approaches to CTC isolation that rely on the reported large size of CTCs compared with WBCs are confounded by the significant overlap between these cell populations, as illustrated here using an enrichment platform that is unbiased by cell size. Established ATCC tumor cell lines are typically larger than WBCs, but primary CTCs in the bloodstream appear to be heterogeneous in size, even within individual patients, with 66% CTCs overlapping in size with 99% of WBCs (**Fig. 4H**). Other non-antibody-based approaches to CTC analysis, such as direct plating of blood cells onto a surface followed by high-speed imaging^41^, cannot be readily scaled up by 100-fold to process the number of cells within a leukopak. Taken together, microfluidic platforms have the advantage of being readily scaled up and automated^7,20^, with the potential to produce a point-of-care device for clinical applications in cancer monitoring.

### Clinical applications of CTC enrichment from leukapheresis samples

Liquid biopsies, primarily centered on DNA mutation analysis from ctDNA, have emerged as powerful tools for monitoring cancer evolution during treatment, including the emergence of mutations that confer resistance to targeted therapeutic drugs^1^. However, the advent of antibody and epitope-specific therapies brings with it a pressing need to measure cell surface protein markers on tumor cells. Most recently, the successful deployment of antibody-drug conjugates, which target cell surface epitopes on cancer cells to deliver chemotherapy payloads, is dependent on the expression of the appropriate cell surface protein^10,11^. Stratifying patients for treatment with the correct ADC will be critical, given the rapidly increasing armamentarium of ADCs targeting different cell surface epitopes, as will on-treatment monitoring for the emergence of resistant cancer cells, some of which may have lost expression of the targeted epitope^42^. Combined with the large number of CTCs harvested by negative depletion of WBCs from leukopaks, the multispectral CTC imaging platform will yield a highly quantitative assessment of epitope expression, without biased selection for EpCAM-positive cancer cells, and it will provide sufficient numbers to identify intra-patient heterogeneity in cell surface marker expression by individual tumor cells. As with all liquid biopsies, CTC measurements also have the advantage of sampling multiple sites of disease rather than a single biopsied lesion, and they can be repeated serially and non-invasively.

In addition to the quantitation of specific cell surface epitopes for immune therapies, the ability to interrogate large numbers of intact cancer cells from the blood in individual patients provides unprecedented opportunities for serial multi-omics at the single-cell level, including paired RNA and DNA analyses applied to large numbers of individual CTCs. In the prototype case of prostate cancer described here, leukopak-derived CTC analysis using WES reproduced the mutations identified by cancer gene mutation panel of a resected metastatic lesion (*PTEN* and *CDH1),* and matched RNA analysis identified two unsuspected divergent cancer cell populations, one driven by FGFR signaling and one with inflammation-associated signatures. Indeed, the development of analytics based on CTC-derived RNA will provide exceptional opportunities to understand heterogeneous subpopulations of cancer cells in advanced, treatment-refractory cancers, as well as to achieve non-invasive pharmacodynamic measurements of drug response through monitoring drug-mediated alterations in transcriptional outputs^28^.

Finally, we note the importance of single-cell CNV analysis in the evaluation of CTCs. While apparently even rarer than CTCs, some non-malignant epithelial cells and reactive stromal cells may be shed along with cancer cells from the tumor micro-environment, and rare endothelial fibroblasts or other non-hematological and non-cancerous cells may be harvested, particularly in high volume blood screening^43–45^. Single-cell CNV provides the most efficient and compelling identification of a tumor cell since aneuploidy is a unique and defining characteristic of cancer, and it is readily measured using low-pass NGS sequencing without requiring prior knowledge or interpretation of point mutations that may be of uncertain significance. In this context, the combination of high throughput CTC screening and definitive CNV measurement may have important applications in the blood-based detection of localized cancer. CTCs are currently detectable within 10-20 mL of whole blood using microfluidic enrichment in approximately 10-20% of patients with localized prostate or breast cancers^28,30,46,47^. While not examined here, patients with invasive localized cancers will likely also have a dramatic increase in CTC collection from leukopaks, and robust CNV determination may prove to be a definitive companion test for emerging ctDNA-based cancer screening methods based on interpretation of altered DNA methylation or fragment sizes.

While leukapheresis, coupled with high-throughput CTC enrichment, presents a strategy for clinical deployment of a whole cell-based liquid biopsy, leukapheresis requires specialized clinical facilities, which imposes a practical hurdle, compared with the collection of a standard 10mL blood tube. The patients with metastatic cancer who underwent leukapheresis for this study did not have procedure-related complications, nor have such been reported in other leukapheresis studies^15,18^. Nonetheless, it is likely that leukapheresis-based CTC analyses will be ideally suited for specific clinical decision points, and the fractional blood volume to be sampled may be reduced in cancer types with high CTC burdens.

## Conclusion

Through the clinical application of a high volume and high throughput microfluidic device, the ^LP^CTC-iChip, we have demonstrated the ability to purify large numbers of CTCs from leukapheresis products in patients with metastatic cancer. The technology is feasible, scalable, and can be developed as a point-of-care technology, ultimately enabling a genuine “cell-based liquid biopsy” with multi-analyte analyses applicable to a diverse array of cancers. Leukapheresis-based CTC analysis may enable many clinical applications that have previously been limited by the small numbers of CTCs present in 10-20 mL of blood and allow a comprehensive approach to liquid biopsies that will help pave the way for more effective and personalized cancer management strategies.

## Acknowledgment

We are grateful to the patients who donated blood to enable this work, and to the clinical and research staff at the Mass General Blood Transfusion Service and Cancer Early Detection and Diagnostics Program. This work was supported by the National Institute of Biomedical Imaging and Bioengineering (P41EB002503), the National Cancer Institute (U01CA214297, R21CA260989, and R01CA255602), and by the Howard Hughes Medical Institute.

