## Supplementary File for "Tumor cell-based liquid biopsy using high-throughput microfluidic enrichment of entire leukapheresis product"

[g] Shriners Children's Boston, Massachusetts, 02114, USA.

### Equal contribution

* Corresponding authors

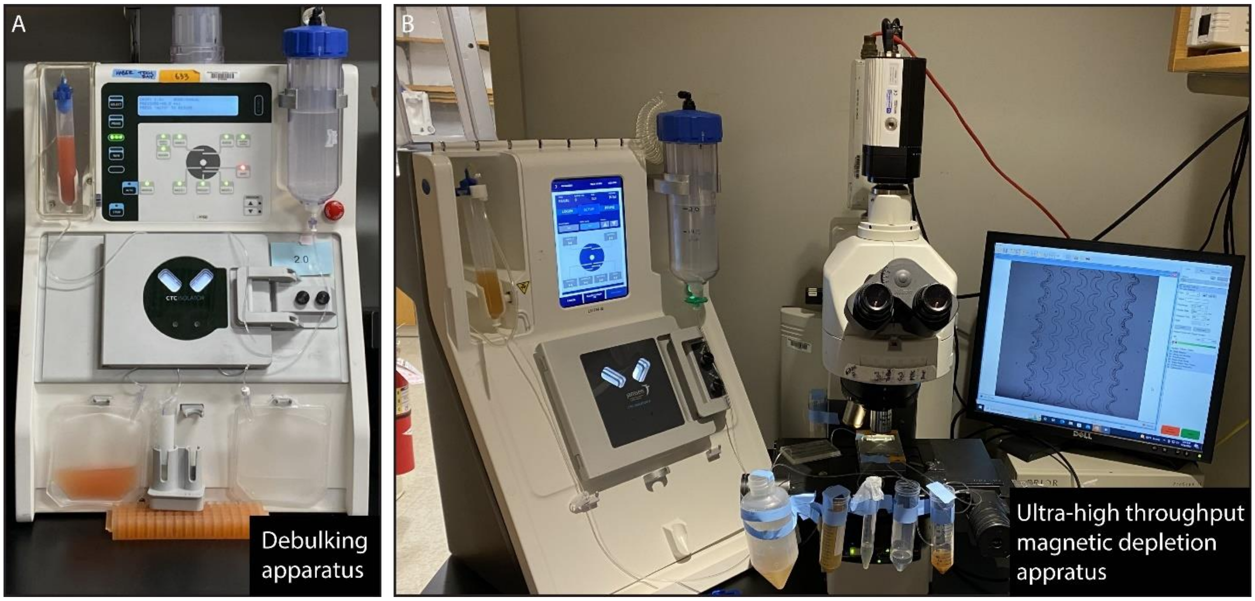


**Figure S1:** The apparatus used for microfluidic isolation of CTCs from leukopaks. (A) The debulking apparatus consists of a pressure source to modulate sample and buffer flow rates. It allows for the collection of WBCs and CTCs from leukopaks into a clean buffer while removing RBCs, platelets, and plasma. (B) The magnetic depletion apparatus uses an aluminum adapter to hold the magnets and the magnetic sorter securely while utilizing the same pressure source as the debulking system. The ultrahigh-throughput magnetic sorter removes magnetically tagged WBCs away from the unmanipulated CTCs.

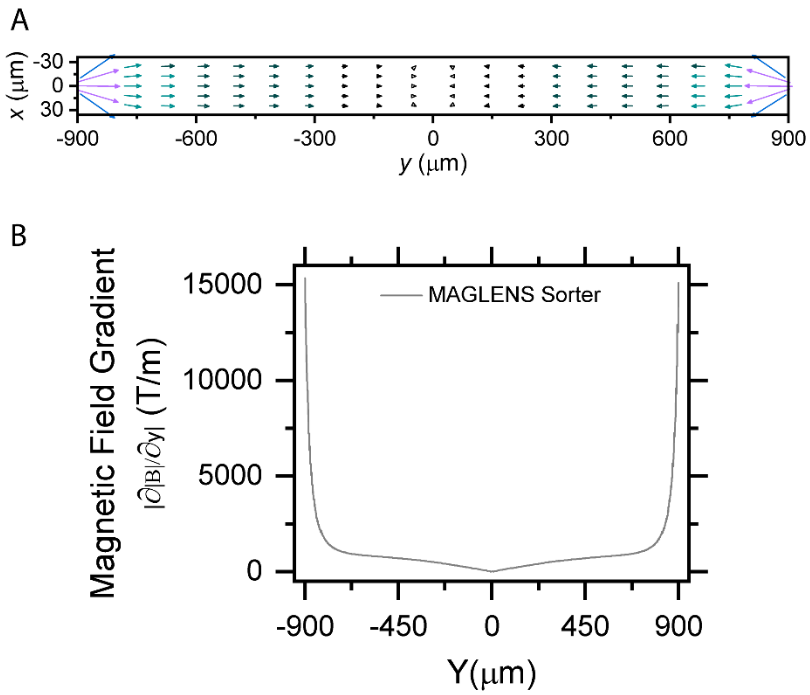


**Figure S2:** (A) Vector plot of magnetic gradient in the sorting channel. Magnetic forces vanish in the center of the channel, creating an inherently safe design for sorting billions of cells without clogging. (B) A high magnetic gradient created by magnetic lenses.

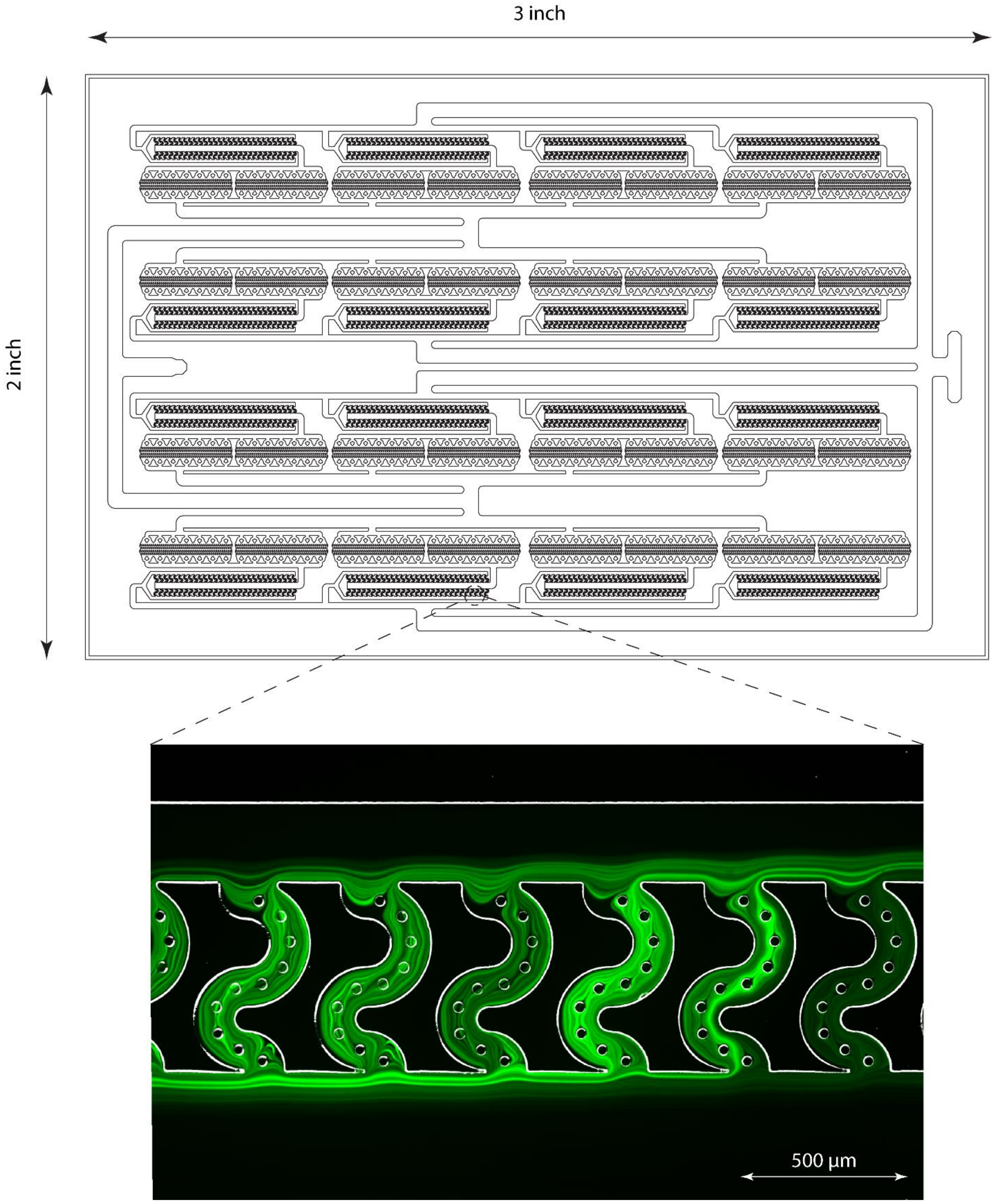


**Figure S3:** Filter chip for removing large clots or aggregates of cells. The inset shows streak images of fluorescently labeled leukocytes flowing through the filter.

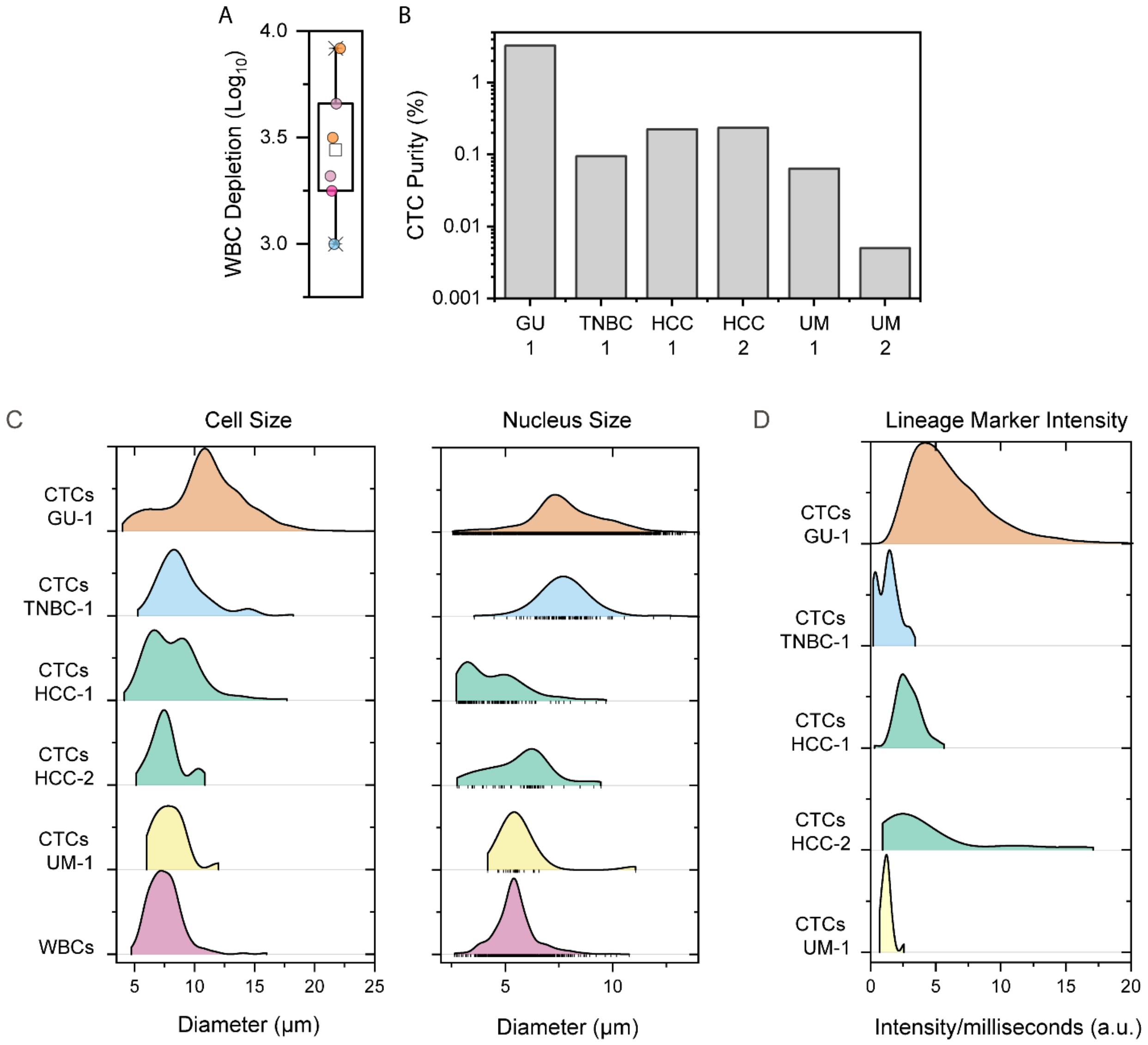


**Figure S4:** (A) WBC depletion after LPCTC-iChip processing of leukopak samples from various cancers. This approach, on average, results in the removal of 99.96% of WBCs while recovering 100 to 58125 CTCs. (B) Bar graph showing CTC purity following enrichment. (C) Measured whole cell and nucleus diameters of individual CTCs, compared with WBCs. (D) Variation across individual CTCs from cases GU-1, TNBC-1, HCC-1, HCC-2, and UM-1 in their intensity of staining for the combined lineage markers.

A

B

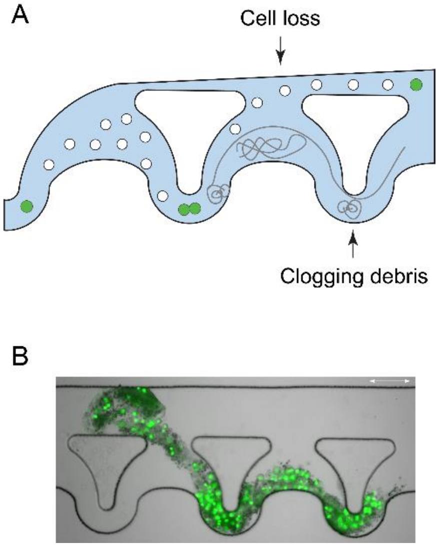


**Figure S5:** Clogging of an inertial concentrator by neutrophil extracellular traps (NETS) in the absence of DNAse. (A) NETS wrap around siphoning pillars and clog the channels, resulting in cell loss. (B) An image of a clogged microfluidic concentrator. Cells are labeled with a fluorescent Dye Cycle Green marker. The scale bar is 100 µm.

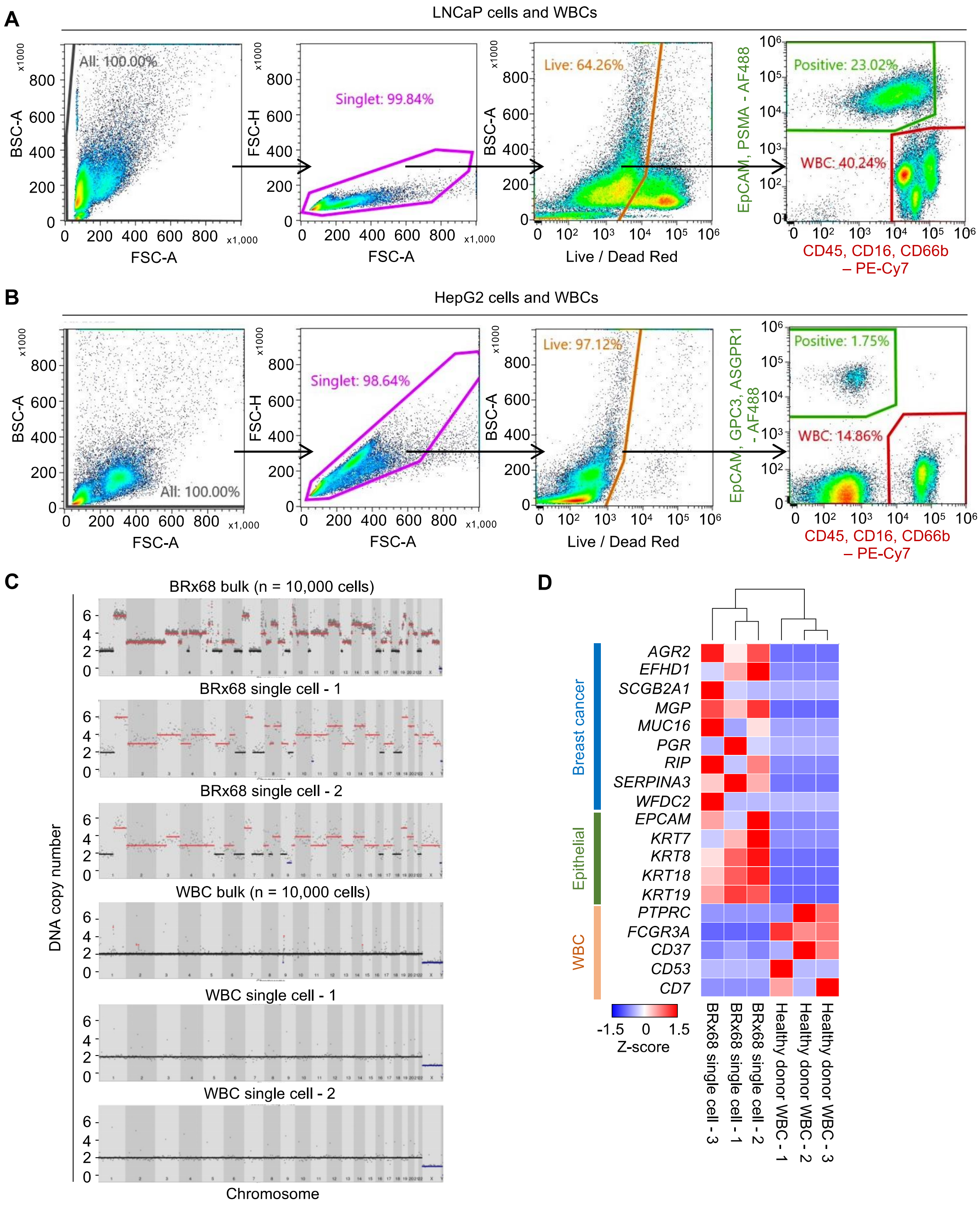


**Figure S6: Development of flow cytometry antibody panels to isolate single CTCs from prostate and liver cancer leukopaks and associated single-cell analysis.** (A) Representative FACS sorting using a customized antibody panel including Alexa fluorophore 488-conjugated antibodies against EpCAM and PSMA and PE-Cy7-conjugated antibodies against WBC markers CD45, CD16, and CD66b. A mixture of cells from the prostate cancer cell line LNCaP and healthy donor WBCs was used to test this panel. (B) Representative FACS sorting using customized antibody panels, including Alexa fluorophore 488-conjugated antibodies against EpCAM, GPC3, and ASGPR1 and PE-Cy7-conjugated antibodies against WBC markers CD45, CD16, and CD66b. A mixture of cells from the liver cancer cell line HepG2 and healthy donor WBCs was used to test this panel. (C) Representative DNA copy-number variation (CNV) analysis in bulk and in single cells, derived from the breast CTC culture BRx68, compared with diploid genomes in bulk and single leukocytes. Ginkgo was used for DNA copy number analysis from whole genome sequencing data. (D) Supervised hierarchical clustering of z-transformed heatmap of gene expression in single-cell RNA-seq data. The cultured BRx68 CTCs have high expression of epithelial and breast cancer lineage markers and absent expression of leukocyte (WBC) markers. Single WBCs that persisted after processing through the microfluidic device are shown as negative controls.

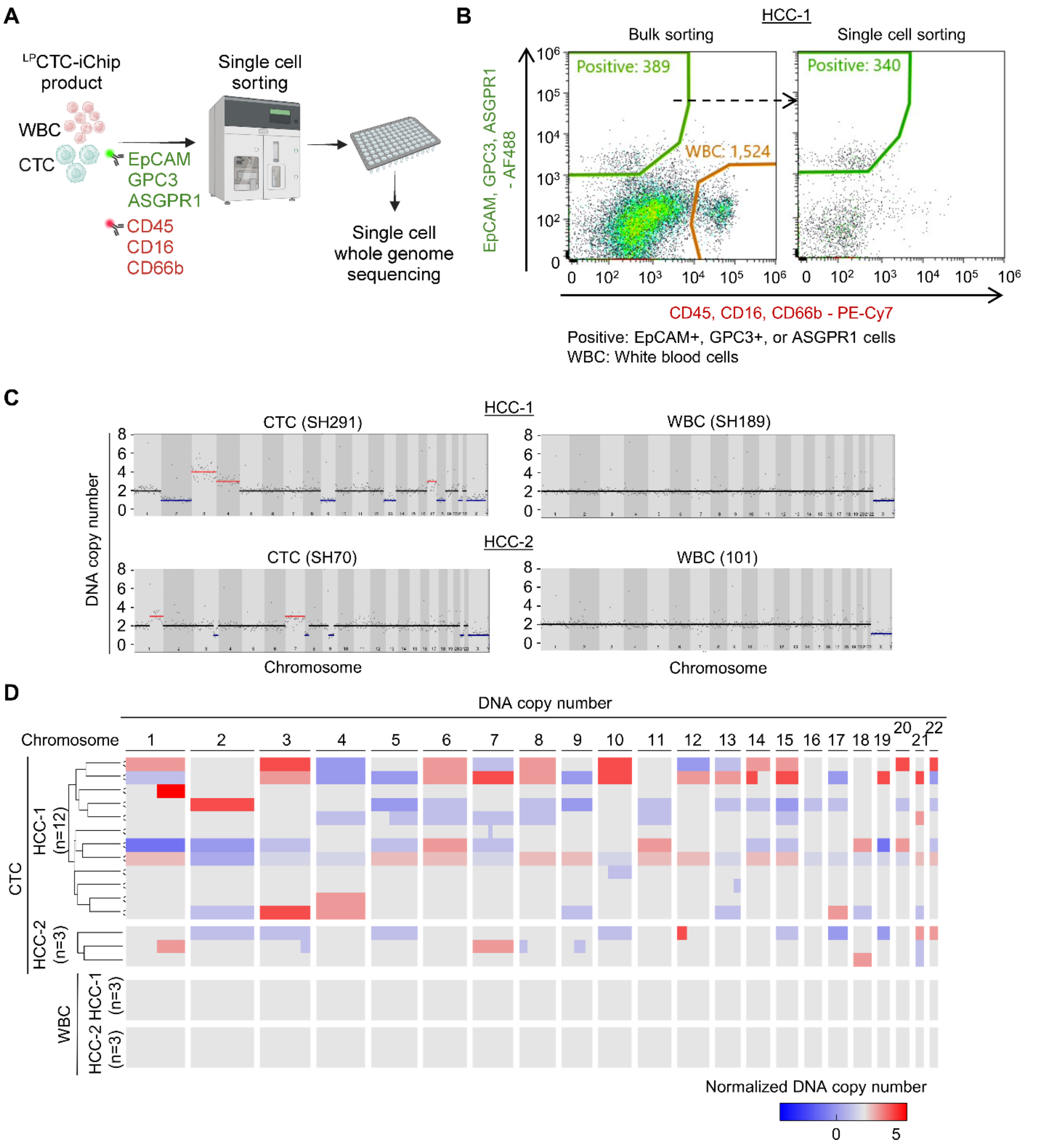


**Figure S7: Isolation of single CTCs from the leukapheresis products of patients with hepatocellular cancer (HCC) and single-cell DNA copy number analysis.** (A) Schematic of single-cell isolation using FACS sorting of LPCTC-iChip-enriched products, derived from leukopak samples of patients with metastatic HCC, followed by single-cell whole genome sequencing. (B) Two-step-sorting strategy using FACS sorting (SONY sorter): initial bulk sorting into a tube to remove dead cells and contaminating WBCs is followed by single-cell sorting to isolate individual CTCs into wells on a plate. Sorting is performed using pooled AF488-conjugated antibodies against the epithelial marker EpCAM and the liver-specific markers GPC3 and ASGPR1 *versus* pooled PE-Cy7-conjugated antibodies against WBC markers CD45, CD16, and CD66b. (C) Representative DNA copy-number variation (CNV) analysis in individual CTCs, compared with diploid WBCs from patients HCC-1 and HCC-2. Ginkgo was used for DNA copy number analysis from single-cell whole genome sequencing data. (D) Unsupervised hierarchical clustering of the normalized heatmap showing individual cell CNVs derived from single-cell whole genome sequencing data of liver CTCs and WBCs. CTCs from HCC-2 show two distinct subpopulations based on CNV analyses. WBCs are shown as negative controls. Ginkgo was used for DNA copy number analysis.

**Table S1:** Patient details and leukapheresis parameters

| Tumor type | Tumor stage | Total blood volume processed (mL) | Duration (minutes) | Machine used | Blood processing flow rate on the apheresis machine (mL/minute) |
|---|---|---|---|---|---|
| GU-01 | Stage IV | 6379 | 109 | Spectra Optia | 60.0 |
| TNBC-01 | Stage IV | 5991 | 120 | Spectra Optia | 51.4 |
| HCC-01 | Stage IV | 4565 | 107 | Spectra Optia | 40.0 |
| HCC-02 | Stage IVB | 7176 | 120 | Spectra Optia | 60.0 |
| UM-1 | Stage IIIC | 6523 | 120 | Spectra Optia | 60.0 |
| UM-2 | Stage IIIA | 4345 | 120 | Spectra Optia | 50/32 |

**Table S2:** List of antibodies for immunofluorescence staining

| **Antibody** | **Vendor** | **Catalog Number** | **Clone** |
|---|---|---|---|
| EpCAM – AF488 | Cell Signaling | 5198S | VU1D9 |
| Pan-Keratin (C11) – AF488 | Cell Signaling | 4523S | |
| Cytokeratin 19 – AF488 | Invitrogen | MA5-18158 | A53-B/A2 |
| CD16 – AF647 | BioLegend | 302020 | 3G8 |
| CD45 – AF647 | BioLegend | 304056 | HI30 |
| CD66b – AF647 | BioLegend | 305110 | G10F5 |
| Sox10 – AF488 | Abcam | 270150 | SP267 |
| Melan-A – AF488 | Abcam | 200544 | EP1422Y |
| NG2/MCSP – AF488 | R&D Systems | FAB2585G | LHM-2 |
| ASGR1 – FITC | Novus Biologicals | NBP1-51109 | 8D7 |
| GPC3 – AF488 | Novus Biologicals | NBP2-47763AF488 | 1G12 + GPC3/863 |
| PSMA – AF488 | Invitrogen | MA5-18161 | GCP-05 |
| CD16 – PE-Cy7 | BioLegend | 980110 | 3G8 |
| CD45 – PE-Cy7 | BioLegend | 982310 | HI30 |
| CD66b – PE-Cy7 | BioLegend | 396910 | QA17A51 |
| LIVE/DEAD™ Fixable Red | Invitrogen | L34971 | |

**Table S3:** List of ddPCR probes

| Test | Primer | Exon Junction | Assay Name | Probe | Primer 2/REV | Primer 1/FWD |
|---|---|---|---|---|---|---|
| Prostate | **FOLH1** | 10-11 | Hs.PT.58.38 521043 | /5HEX/ATG AAC AAC/ZEN/AGC TGC TCC ACT CTG A/3IABkFQ/ | CAATGT GATAGG TACTCTC AGAGG | TGTTCCA AAGCTC CTCACA A |
| Prostate | **KLK2** | 2a-3 | Hs.PT.58.20 108543 | /56-FAM/TGG CTA TTC/ZEN/TTC TTT AGG CAA TGG GCA/3IABkFQ/ | GCTGTG TACAGT CATGGA TGG | GTCTTCA GGCTCA AACAGG T |
| Prostate | **KLK3** | 5 | Custom-Designed | /56-FAM/TGT GCT TCA/ZEN/AGG TAT CAC GTC ATG GG/3IABkFQ/ | CTTTCG GGCAGG GCACAT | GGGCCC ACTTGTC TGTAATG G |
| Prostate | **TMPRSS2** | 12-13 | Hs.PT.58.22 365438 | /56-FAM/ACC CGG AAA/ZEN/TCC AGC AGA GCT/3IABkFQ/ | CCCAAC CCAGGC ATGATG | TCAATGA GAAGCA CCTTGG C |
| Prostate | **FAT1** | 13-14 | Hs.PT.58.45 775110 | /56-FAM/TCT TGT CAG/ZEN/CAG CGT TCC CGG/3IABkFQ/ | GATCCTT ATGCCA TCACCG T | ATCAGC AGAGTC AATCAGT GAG |
| Prostate | **STEAP2** | 4-5 | Hs.PT.58.19 467586 | /5HEX/ACA TGG CTT/ZEN/ATC AGC AGG TTC ATG CA/3IABkFQ/ | CATGTT GCCTAC AGCCTC T | TCTCCAA ACTTCTT CCTCATT CC |
| Prostate | **AGR2** | 7-8 | Hs.PT.58.38 683802 | /56-FAM/ATG CTT ACG/ZEN/AAC CTG CAG ATA | CTGACA GTTAGA GCCGAT ATCAC | CAATTCA GTCTTCA GCAACT TGAG |

| | | | | | | |
|---|---|---|---|---|---|---|
| | | | | CAG CTC/3IABkFQ/ | | |
| Prostate | **HOXB13** | 1-2 | Hs.PT.58.4375164 | /5HEX/CAG CAT TTG/ZEN/CAG ACT CCA GCG G/3IABkFQ/ | CAGCCAGATGTGTTGCCA | CTGTACGGAATGCGTTTCTTG |
| Prostate | **AR-wt** | 4-5 | Custom-Designed | /56-FAM/ACG ACC AGA/ZEN/TGG CTG TCA TTC AGT/3IABkFQ/ | CCAGCCCATGGCAAACA | CTGGGAGAGAGACAGCTTGTA |
| Prostate | **AR-V7** | 3-4 | Custom-Designed | /5HEX/AAGCAGGGA/ZEN/TGACTCTGGGAGAAA/3IABkFQ/ | CTTTCTTCAGGGTCTGGTCATT | CTTGTCGTCTTCGGAAATGTTATG |
| Prostate | **TACSTD2** | 1 | Custom-Designed | /56-FAM/TCG GTC CAA/ZEN/CAA CAG GAA ACC TGA/3IABkFQ/ | CTATGCCATCCCTTCCTTCAC | CAGGGTCTCCTTTCTTTCTCAC |
| Prostate | **MYC** | 2-3 | Custom-Designed | /56-FAM/TTG TTC CTC/ZEN/CTC AGA GTC GCT GC/3IABkFQ/ | CAACATCGATTTCTTCCTCATCTTC | TTCTCTCCGTCCTCGGATT |
| Prostate | **POU3F2** | 1 | Custom-Designed | /56-FAM/TGA AGC TAT/ZEN/CCA GAG CAG GGC AAA/3IABkFQ/ | GAACTCCAGCTTCTGACCTTAC | GGTATGGGAACTGGCCTTTAG |
| Prostate | **EZH2** | 17-18 | Custom-Designed | /5HEX/TTACAGATA/ZEN/CAGCCAGGCTGATGCC/3IABkFQ/ | TCGATGCCGACATACTTCAG | CATCCAGACTGGCGAAGAG |
| Prostate | **SYP** | 6-7 | Custom-Designed | /5HEX/TAG TCT GGT/ZEN/CAG TGA AGC CCA GGA/3IABkFQ/ | GGGTGGAGACCTAGGGTATAG | CACCTCCTTCTCCAATCAGATG |

| | | | | | | |
|---|---|---|---|---|---|---|
| Prostate | **CSPG4** | 9-10 | Custom-Designed | /56-FAM/AGC CAC CTC/ZEN/TGG AAG AAC AAA GGT/3IABkFQ/ | GCCAAG AGATTG GAGGCA T | TGCTGT GGCTGT GTCTTT |
| Prostate | **DLL3** | 7-8 | Custom-Designed | /5HEX/ATG GTC CGA/ZEN/GCT CGT CCG TAG ATT/3IABkFQ/ | GTCTAC ATCTTCA GGGCGA TT | CAACCT AAGGAC GCAGGA G |
| Prostate | **E2F1** | 5-6 | Custom-Designed | /5HEX/AAA CAT CGA/ZEN/TCG GGC CTT GTT TGC/3IABkFQ/ | TGATCC CACCTA CGGTCT C | GGACTC TTCGGA GAACTTT CAG |
| Prostate | **FOXA2** | 3 | Custom-Designed | /5HEX/ACC CGT TCT/ZEN/CCA TCA ACA ACC TCA/3IABkFQ/ | AGTGCA TCACCT GTTCGT AGGC | GGAACA CCACTA CGCCTT CAAC |
| Prostate | **CHGA** | 7-8 | Custom-Designed | /5HEX/AGGAGA AGA/ZEN/AAGA GGAGGAGGGC A/3IABkFQ/ | CTGGTG GGCCAC TTTCTC | TCCAGG TCCGAG GCTAC |
| Melanoma | **TYRP1** | 4-5 | Hs.PT.58.21 339707 | /56-FAM/CTC AAT GGC /ZEN/GAG TGG TCT GTG ACT /3IABkFQ/ | CTC TAA TAA GCC CAA ACT CTG TCT | AGC TGG ATT TCT CCT AAT TGG C |
| Melanoma | **CSPG4** | 1-2 | Hs.PT.58.21 69197 | /56-FAM/CGC GGC TTC /ZEN/CTT CTT CGG TGA /3IABkFQ/ | TTG GCT TTG ACC CTG ACT ATG | GCA GGT CTA TGT CGG TCA G |
| Melanoma | **MAGE A1** | 1-3 | Hs.PT.58.27 428541 | /56-FAM/CTG TCC TCT /ZEN/GGG TTG GCC TGT /3IABkFQ/ | GCG AGG TTT CCA TTC TGA GG | CAG ATC TTC TCC TTG GTG CTC |
| Melanoma | **GAGE1** | 4-6 | Hs.PT.58.25 043287 | /56-FAM/AGG CGT TTT /ZEN/CAC CTC | GTG TGA AGA TGG | CCT GCA ACA TTT |

| | | | | | | |
|---|---|---|---|---|---|---|
| | | | | CTC TGG ATT T/3IABkFQ/ | TCC TGA TGG | CAG CAT GTC |
| Melanoma | **MAGE A4** | 1-3 | Hs.PT.58.27 894043 | /56-FAM/CCA CAG GCA /ZEN/GAT CTT CTC CTT GGT G/3IABkFQ/ | CTG TGA GGA GTC AAG GTT CTG | CAA GAG TGC AGG CAA AAG C |
| Melanoma | **TNC** | 26-27 | Hs.PT.58.39 402235 | /56-FAM/CAC CTG CTG /ZEN/TCC CAC TGT ACC C/3IABkFQ/ | GAC TCG CTA CAA GCT GAA GG | GTT GGT GAT GGC TGA ATC TGT |
| Melanoma | **IL13RA 2** | 8-9 | Hs.PT.58.40 296980 | /56-FAM/ACC TTC CCA /ZEN/GCA TTG TTT ATC ACT CCA /3IABkFQ/ | CAA ATG AAA CCC GAC AAT TAT GC | GTA GCC AGA AAC GTA GCA AAG |
| Melanoma | **MAGE A2** | 3-5 | Hs.PT.58.23 66134 | /56-FAM/TCG GGT CCT /ZEN/ACT TGT CAG CCT GT/3IABkFQ/ | GAT GCA GTG GTT CTA GGA TCT G | GAT CTT CTC CTT CAA TGC TCC T |
| Melanoma | **MAGE C2** | 1-3 | Hs.PT.58.23 276508 | /56-FAM/CTG TGC TGA /ZEN/CTT TAG GCT GTG TGC /3IABkFQ/ | ACC TCC CAC CAT AGA GAG AAG | CCT TGA CTC CTG GCA CTG |
| Breast | **CXCL1 4** | 2-3 | Hs.PT.58.19 273291 | | | |
| Breast | **FAT1** | 13-14 | Hs.PT.58.45 775110 | /56-FAM/TCT TGT CAG /ZEN/CAG CGT TCC CGG /3IABkFQ/ | GAT CCT TAT GCC ATC ACC GT | ATC AGC AGA GTC AAT CAG TGA G |
| Breast | **FAT2** | 9-10 | Hs.PT.58.24 846942 | /56-FAM/ACC TGC TAC /ZEN/ATC ACA GAG GGA GAC C/3IABkFQ/ | CCT GGA TGC TGA CAT TTC TGA | TCC TCC ACT CAT CTC CAA CT |
| Breast | **MGP** | 2-4 | HS.PT.58.63 5768 | | | |

| | | | | | | |
|---|---|---|---|---|---|---|
| Breast | **MUC16** | 5-6 | Hs.PT.58.35 43722 | | | |
| Breast | **PRAM E** | 3-4 | Hs.PT.58.45 281469 | /56-FAM/CAA GCG TTG /ZEN/GAG GTC CTG AGG C/3IABkFQ/ | GCA ACA AGT GAC TGA GAC CTA | GTC CAC ACA CTC ATG CTG AT |
| Breast | **SERPI NA3** | 4-5 | Hs.PT.58.15 580605 | | | |
| Breast | **SFRP1** | 1-3 | Hs.PT.58.38 429156 | /56-FAM/TGT GAC AAC /ZEN/GAG TTG AAA TCT GAG GCC /3IABkFQ/ | CAA TGC CAC CGA AGC CT | CTT TTA TTT TCA TCC TCA GTG CAA AC |
| Breast | **TMPRS S4** | 9-11 | Hs.PT.58.31 61735 | | | |
| Breast | **CXCL1 3** | 2-3 | Hs.PT.58.45 801487 | | | |
| Breast | **PGR** | 7-8 | HS.PT.58.15 66542 | | | |
| Breast | **SFRP2** | 1-2 | Hs.PT.58.20 705989 | | | |
| Breast | **AGR2** | 7-8 | Hs.PT.58.38 683802 | | | |
| Breast | **EFHD1** | 4-5 | Hs.PT.58.27 534728 | | | |
| Breast | **PIP** | 2-3 | Hs.PT.58.19 165954 | | | |
| Breast | **SCGB2 A1** | 1-2 | Hs.PT.58.86 4035 | | | |
| Breast | **WFDC 2** | 2-3 | Hs.PT.58.25 117187 | | | |

**Table S4:** Unsupervised gene set enrichment analysis for single-cell RNA-seq data of prostate CTC with CNVs.

**Table S5:** Whole exome sequencing revealed variants of clinically unknown significance in prostate CTCs.

#### Materials and Methods

##### Microfabrication

The Magnetic Sorter chips were designed and produced at Massachusetts General Hospital (MGH) using soft lithography with Polydimethylsiloxane (PDMS). In a nutshell, this involved applying a layer of SU-8 50 at 2200 rpm for a duration of 30 seconds onto a silicon wafer that had been pre-baked. Photolithography was then utilized to pattern channels by exposing the SU-8 layer to 365 nm UV radiation through a mylar mask. The exposed silicon wafer was subsequently processed with Baker BTS-220 SU-8 developer to form microfeatures.

To create a 1-mm thick PDMS layer, a Sylgard 184 PDMS kit was mixed in a 7:1 ratio of base to cross-linker and poured onto the SU-8 mold. After degassing within a vacuum chamber, the PDMS layer was cured in a convection oven at 65°C for a period of 12 hours. Given the device's thickness of only 1 mm, inlet and outlet ports for press-fit tubing connections were established using 3 mm thick rectangular PDMS pads. These thicker PDMS pads were generated by depositing PDMS onto a featureless SU-8 50 mold in a 10:1 ratio of base to cross-linker. The attachment of these rectangular pads to the device was achieved via plasma bonding, and holes were punctured using a 1.2 mm biopsy punch. Subsequently, the device was bonded to a 3-inch × 1.5-inch glass slide that had been cleaned with piranha solution, followed by baking at 85°C for 10 minutes. This was succeeded by an extended bake for 3 hours at 150°C to augment the bonding and Young's modulus of the PDMS.

To house the magnetic sorter chip, a manifold was designed, accommodating six permanent magnets arranged in two parallel quadrupolar configurations. This manifold was machined from 6061 aluminum alloy, with screws securing the magnets in place. For more comprehensive information on PDMS fabrication and the magnetic manifold design, we encourage you to consult our previous report[1].

Debulking chips were fabricated with medical-grade cyclic olefin copolymer (COC) using variotherm injection molding by Stratec Biomedical AG (Germany). Each debulking device included 16 inertial separation array devices.

##### Apheresis Collection

Leukapheresis on cancer patients was performed at the MGH Apheresis Center using an experimental protocol reviewed and authorized by the MGB Institutional Review Board (MGB IRB 2020P000251). Leukopaks were collected on approximately one human blood volume using the Spectra Optia system in the continuous mononuclear cell collection mode. All five patients tolerated apheresis well, and no adverse event was reported. On average, the apheresis took 2 hours to complete. Whole blood donation from healthy donors at our lab was collected using MGB IRB Protocol # 2009-P-000295. In some cases, whole blood samples from healthy donors were also procured from Research Blood Components, LLC (Brighton, MA).

##### WBC depletion antibody preparation

The WBC depletion cocktail included three antibodies: biotinylated anti-human CD45 (Thermo Fisher Scientific, clone HI30, IgG1, 0.25 µg/million cells), biotinylated anti-human CD16 (BD Biosciences, clone 3G8, IgG1, 0.05 µg/million cells), and biotinylated anti-human CD66b (Novus Biologicals, clone 80H3, IgG1, 0.025 µg/million cells). This antibody cocktail was prepared by mixing antibodies in sterile filtered 1X phosphate-buffered saline (PBS) and 0.1% BSA and stored at 4°C for use within 7 days.

**Magnetic beads preparation**

To label WBCs, we employed 1 µm Dynabeads MyOne Streptavidin T1 superparamagnetic beads from Invitrogen. The beads underwent three rounds of magnetic washing using 0.01% Tween 20 in 1X PBS, followed by three additional washes with 0.1% BSA in 1X PBS to minimize nonspecific binding. Subsequently, these streptavidin-conjugated beads were stored at 4°C in a 0.1% BSA solution at 10 mg/mL. Washed beads were utilized within a week.

**Single-cell sorting:** The enriched leukopak by $^{LP}$CTC-iChip were Fc blocked (Jackson ImmunoResearch Laboratories, Cat# 009-000-008) and then immunostained with AF488-conjugated EpCAM (Cell signaling, Cat# 5198) / PSMA (Invitrogen, Cat# MA5-18161) / GPC3 (Novus Biologicals, Cat# NBP2-47763AF488) / FITC-conjugated ASGPR1 (Novus Biologicals, Cat# NBP1-51109) antibodies and PE-Cy7-conjugated CD45 (Biolegend, Cat# 982310) / CD16 (Biolegend, Cat# 980110) / CD66b (Biolegend, Cat# 396910) antibodies, and LIVE/DEAD Red (Invitrogen, Cat# L34971). Cells were treated with DNase I (Worthington Biochemical Corporation, Cat# LS006361) to reduce cell clumping in single-cell suspensions before cell sorting. We performed two-step sorting methods (the ultra-yield bulk sorting followed by the single cell plate sorting) to isolate single viable CTCs (EpCAM or PSMA positive for prostate cancer and EpCAM or GPC3 or ASGPR1 positive for liver cancer) and WBCs (CD45/CD16/CD66b positive) using SONY sorter SH800. The single viable cells were individually sorted into 96-well PCR plates containing the cell lysis buffer for downstream single-cell whole genome sequencing and RNA sequencing.

**Paired single-cell WGS and RNA-seq:** For these experiments, we used either single CTCs or WBCs individually sorted by SONY sorter from $^{LP}$CTC-iChip enriched leukopak. These were subjected to single-cell whole genome sequencing and RNA-seq analysis to obtain DNA copy number and gene expression at the single-cell resolution. Briefly, single cells were first lysed in 5 µl of single-cell lysis buffer, and 0.5 µl of Magnetic MyOne Carboxylic Acid Beads (Invitrogen, Cat# 65011) were then added to each single cell lysate to facilitate segregation of nucleus versus cytoplasm[2]. After magnetic separation, the pellet (aggregated beads with the intact nucleus) was resuspended in 5 µl of DNA lysis buffer and subjected to single-cell whole genome amplification using the MALBAC protocol[3], while the supernatant containing cytoplasmic RNA was converted to cDNA and amplified using Smart-seq2 protocol[4]. The sequencing libraries were constructed using Nextera XT DNA library preparation kit (illumine, Cat# FC-131-1096) and sequenced on the Illumina NovaSeq 6000 platform with 150 bp paired-end reads and 1.5-2x genome coverage.

**WGS pipeline:** Reads were trimmed using Trim Galore! (https://www.bioinformatics.babraham.ac.uk/projects/trim_galore/) version 0.4.3 with default settings. Reads were then aligned to version hg19 of the human genome using version 0.7.15 of bwa mem[5] with default settings. We applied the MarkDuplicates function of the Picard library (http://broadinstitute.github.io/picard) and then discarded from the resulting BAM file any reads for which the UNMAP, SECONDARY, QCFAIL, DUP, or SUPPLEMENTARY flags were set. We then used the bamToBed program of version 2.18.2 of bedtools and uploaded the resulting bed files to the Ginkgo website (http://qb.cshl.edu/ginkgo). We processed the samples using

Ginkgo's default settings, except "mapped with" was set to "bwa," and for single cells, "bin size of" was set to 5Mb.

**Copy number heatmap:** We used the copy number values in the SegCopy spreadsheets that can be downloaded from the Ginkgo website. To compensate for Ginkgo's tendency to sometimes assign copy numbers that are too high, we subtracted the median copy number from the copy number for each sample. Then we added 2 to form a Normalized DNA copy number. To cluster the Normalized DNA copy number profiles, we used hierarchical clustering as implemented by the hclust function of version 3.6.3 of R with the average agglomeration method and distance defined by the Pearson correlation coefficient.

**Variability score:** We computed a variability score (VS) as described in the supporting information for Knouse et al.[6] with the exceptions that we used 1-Mb bins instead of 500-kb bins and used median absolute deviation (MAD) instead of SD and used median instead of average. We considered samples with this VS greater than 0.85 to be of low quality.

**RNA-seq pipeline:** Raw fastq reads generated from the HiSeq X sequencer were first cleaned using TrimGalore (v0.4.3) (https://github.com/FelixKrueger/TrimGalore) to remove the adapter-polluted reads and reads with low sequencing quality. Cleaned reads were aligned to the human genome (hg19) using Tophat (v2.1.1)[7]. PCR duplicates were further removed using samtools (v1.3.1)[8], and gene counts were computed using HTseq (v0.6.1)[9]. To cluster the RNA-seq profiles we used the 2,000 genes with the highest standard deviation across the samples. We then used hierarchical clustering as implemented by the hclust function of version 3.6.3 of R with the ward.D agglomeration method and distance defined by Pearson correlation coefficient.

**Gene Set Enrichment Analysis (GSEA)**: We first removed genes with 0.9 quantile of reads-per-million (RPM) less than 10. We then ran version 2.0.14 of the UCSD/BROAD gsea2 software[10] in preRanked mode on the BIOCARTA, GO, HALLMARK, KEGG, MIR, PID, REACTOME, and TFT gene sets from version 7.1 of the MSigDB (https://www.gsea-msigdb.org/gsea/msigdb/). The preRanked values were sign(logFoldChange) times-log10 of the var.equal t-test p-value for the two classes being compared.

**WES and mutation analysis:** We pooled MALBAC-amplified genomic DNA from 26 CTCs displaying clear CNV as the pseudo-bulk DNA. Exomes were captured using Agilent SureSelect Human All Exon V6 Kit and sequenced on the Illumina NovaSeq 6000 platform with 150 bp paired-end reads and 100x coverage. We ran the pipeline described above for WGS. We then ran MuTect on the resulting BAM to detect mutations and employed the following four filters: (1) the mutation must not be found in a panel of normals, (2) the mutation must be in a gene that is in the Cancer Gene Census (https://www.sanger.ac.uk/data/cancer-gene-census), (3) the mutation must be protein-altering, i.e., coding nonsynonymous, and (4) the mutation must have an allele frequency of at least 30% and be supported by at least 3 reads. This yielded 48 mutations.

**Cytospin and immunostaining**

For immunofluorescence-based CTC enumeration, a part of $^{LP}$CTC-iChip product was fixed using 0.5% PFA for 10 minutes, followed by cytospin at 2000 rpm for 5 minutes using the Thermo Scientific Cytospin 4 centrifuge and Epredia EZ Megafunnel. After cytospin, slides were washed using cold 1X-PBS, followed by nonspecific blocking for 1 hour with 3% BSA and 2% Normal Goat Serum solution. This was followed by staining for 1 hour with primary antibodies for nucleus markers, tumor markers, and leukocyte markers, followed by washing with 1X-PBS. Please refer to Table S2 for a complete list of antibodies used in this work. Slides were subsequently coverslipped using a mounting medium (ProLong Diamond Antifade Mountant with DAPI from Life Technologies) and left to cure for 12 to 16 hours before imaging.

##### CTC enumeration method

Prepared slides were scanned using an Akoya Vectra Polaris™ Automated Quantitative Pathology Imaging System. High-content multispectral images were taken at 40X magnification in DAPI, AF488, and AF647 channels, with exposure times in each channel determined from prior spiked cells, healthy donor and whole blood patient samples. These immunofluorescence images were captured using an 8-bit camera, which divides pixel intensity from 0 to 255. The images were subsequently exported and processed using a HALO image analysis platform. First, automated analysis of the entire slide was used to identify individual cells and the levels of fluorescent signal in each channel. Adhering to preset inclusion and exclusion parameters, digital image processing-based segmentation algorithms marked a subset of these cells for further manual verification. These cells were then individually analyzed, considering overall morphology, size, and staining intensity. Following independent analysis by at least two researchers, CTCs were scored and quantified.

##### Droplet Digital PCR analysis

Droplet Digital PCR (ddPCR) to detect the expression of published cancer-specific gene marker panels[11–14] was performed on leukopak-derived RNA. For the hepatocellular carcinoma and breast patient samples, whole transcriptome amplification (WTA) was performed on RNA extracted from 0.5% of the leukopak product using the SMARTer Ultra Low-input RNA kit, version 4 (Takara). During WTA, 18 cycles of amplification were performed. For the uveal melanoma patient samples and prostate cancer patient samples, reverse transcription was performed on RNA extracted from 0.5% of the leukopak product using the SuperScript III First-Strand Synthesis System (Life Technologies). Gene-specific nested PCR amplification was employed for 10 or 14 PCR cycles, depending on the gene, for the uveal melanoma patient samples before ddPCR. No amplification was performed for prostate cancer patient samples. Samples were prepared for ddPCR following the standard protocol (Biorad). Briefly, 2x ddPCR Supermix for probes (no dUTP) was combined with 1-2% of the patient cDNA and 1x primer/probe mix (IDT). After droplet generation, samples were thermocycled for 45 cycles with an annealing temperature of 70°C and an extension temperature of 53°C. As a negative control, healthy donor blood was also analyzed on the leukopak. As a positive control for assay function, 0.1% of cancer cell line RNA was spiked into healthy donor RNA prior to PCR amplification and downstream processing.

##### ddPCR Data Analysis

The number of cDNA copies per reaction, as calculated by the BioRad QuantaSoft or QX Manager software, was used to generate heatmaps. Briefly, the signal was normalized within each marker such that the maximum value was 1 (red). For the UM and TNBC patients, a threshold was drawn at two standard deviations higher than the median healthy donor background signal for a marker, and this value was subtracted from the marker value for all samples. The subtracted value for each marker is adjusted to 0 when the result is negative.
